# Reconstitution of actomyosin networks in cell-sized liposomes reveals distinct mechanical roles of cytoskeletal organization in membrane shape remodeling

**DOI:** 10.1101/2025.05.18.654456

**Authors:** Makito Miyazaki, Fahmida Sultana Laboni, Taeyoon Kim

## Abstract

The actin cortex, a thin layer of actomyosin network beneath the plasma membrane, regulates various cell functions by generating active forces and inducing membrane deformations, including blebs. Although upstream signaling is involved in regulating cell shape, the extent to which downstream actomyosin molecules can control the shape remains elusive. Here, using a minimal reconstituted system with a combination of agent-based computational model, we show that while actin-membrane coupling strength determines the magnitude of membrane deformation, its balance with actin network connectivity governs the bleb initiation mechanism, either by detachment of the cortex from the membrane or by rupture of the cortex. This balance also regulates whether single or multiple blebs form. Furthermore, both experiments and simulations suggest that not only the dense cortical network but also the sparse volume-spanning network actively contributes to regulating bleb number. These findings provide mechanical insights into how cells tune actin network organization to control their shape.

## INTRODUCTION

Cells assemble diverse structures of actin networks to operate biological functions. Among these structures, one of the most ubiquitous structures is the actin cortex, a thin layer of an actomyosin network formed beneath the plasma membrane^1^. The actin cortex is not only a passive scaffold that stabilizes the cell shape, but also plays pivotal roles in regulating various biological functions, such as migration, division, and polarity establishment, by generating active contractile forces ^1,2,3,4^. These forces drive morphological transitions of cells, including the formation of spherical membrane protrusions called blebs. During apoptosis, cells form transient multiple blebs in random directions^5^. By contrast, in many other processes, the number and position of blebs are tightly regulated. During cell migration mediated by the actin cortex, primordial germ cells create a single bleb with a size comparable to the cell body at the front end to drive migration^6,7,8^. Similarly, during amoeboid migration under physical confinement, various epithelial and mesenchymal cells such as HeLa cells also form a single bleb^9^. In contrast, Walker 293 carcinosarcoma cells^10,11^ and *Dictyostelium* cells^12,13^ create relatively small multiple blebs at the leading edge. During cell division, cells form multiple blebs near the poles, possibly to relieve intracellular pressure elevated by furrow ingression and ensure successful cytokinesis^14^.

How can the same key molecular elements of the actin cortex, namely actin filaments (F-actins) and myosin motors, induce various morphological transitions with tunable properties? This remains an open question. Although the upstream biochemical signaling pathways mediated by Rho-family GTPases and phosphoinositides are involved in the regulations^3,5,15^, downstream proteins in the actin cortex directly mediate the morphological transition. Thus, it is important to ascertain the extent to which physical aspects of the actin cortex can control the cell shape. One of the promising but challenging strategies is a bottom-up approach: reconstituting the process from a minimal set of proteins and cell-sized liposomes^16,17^. It has been demonstrated that liposomes containing actin and myosin exhibit different patterns of contraction depending on the actin-membrane interaction strength^18^, and that the addition of anillin induces detectable membrane deformations including blebs^19^. However, these studies primarily focused on post-contraction steady-states and lacked quantitative time-lapse analysis. Recent studies reported bleb formation upon laser ablation of the cortical actin network^20^ or myosin activation by light-induced degradation of a myosin inhibitor^21^. However, these systems could not reconstitute spontaneous bleb formation. As an alternative approach, agent-based computational models have been employed. To recapitulate bleb formation, cells were often simplified as structures consisting of a membrane, actin cortex, and cytoplasm with internal cytoskeleton^23,24,25^. However, these models could not reproduce large bleb formation because it did not account for membrane fluidity, which permits movement of anchor points between the membrane and the cortex. Although some models have incorporated membrane fluidity^26,27,^ the cytoskeletal networks were drastically simplified, limiting the capability and predictability.

Here, we investigate the physical mechanism of morphological transition driven by the actin cortex, combining a minimal reconstituted model with an agent-based computational model. Our reconstituted system, consisting of purified cytoskeletal proteins and cell-sized liposomes, enables independent control of i) actin-membrane coupling strength, ii) actin network connectivity, and iii) actin network distribution inside liposomes. Unlike previous reconstituted systems, myosin activities are temporally regulated by myosin kinase, mimicking regulation in living cells. The agent-based computational model, consisting of an actomyosin network and a deformable membrane, enables molecular-scale spatiotemporal analysis, providing mechanical insights inaccessible to experiments. Using these advantages, we systematically investigate how each parameter affects the morphological transition process.

## RESULTS

### Reconstituting the actin cortex inside a cell-sized liposome

We developed a minimal model of the actin cortex by reconstituting a contractile actomyosin network from purified proteins (**Fig. S1a**) inside a cell-sized liposome (**Fig. 1a**). In cells, myosin contractility is regulated by phosphorylation of the myosin regulatory light chain by myosin kinases^28^. Phosphorylated myosin dimers assemble into submicrometer-long filaments that generate contractile forces on F-actin^28^ (**Fig. 1a**, inset). To recapitulate this regulation, we used ZIP kinase^29^ (ZIPK), one of the major myosin kinases involved in cell motility^30^, cytokinesis^31,32^, and apoptosis^33^. We tuned the ZIPK concentration to let the cortex formation and contraction processes take place sequentially (**Fig. 1b, Movie S1**). We confirmed by biochemical assays^34,35^ (**Fig. 1c,d**) that 90% of 10 μM actin was polymerized ∼10 min after the encapsulation (**Fig. 1c**, green line), whereas it took ∼40 min to phosphorylate 90% of 1 μM myosin (**Fig. 1c**, magenta line). We also confirmed by fluorescence microscopy that myosin self-assembled into submicrometer-long filaments upon its phosphorylation (**Fig. 1e, Fig. S1b**).

**Figure 1.**
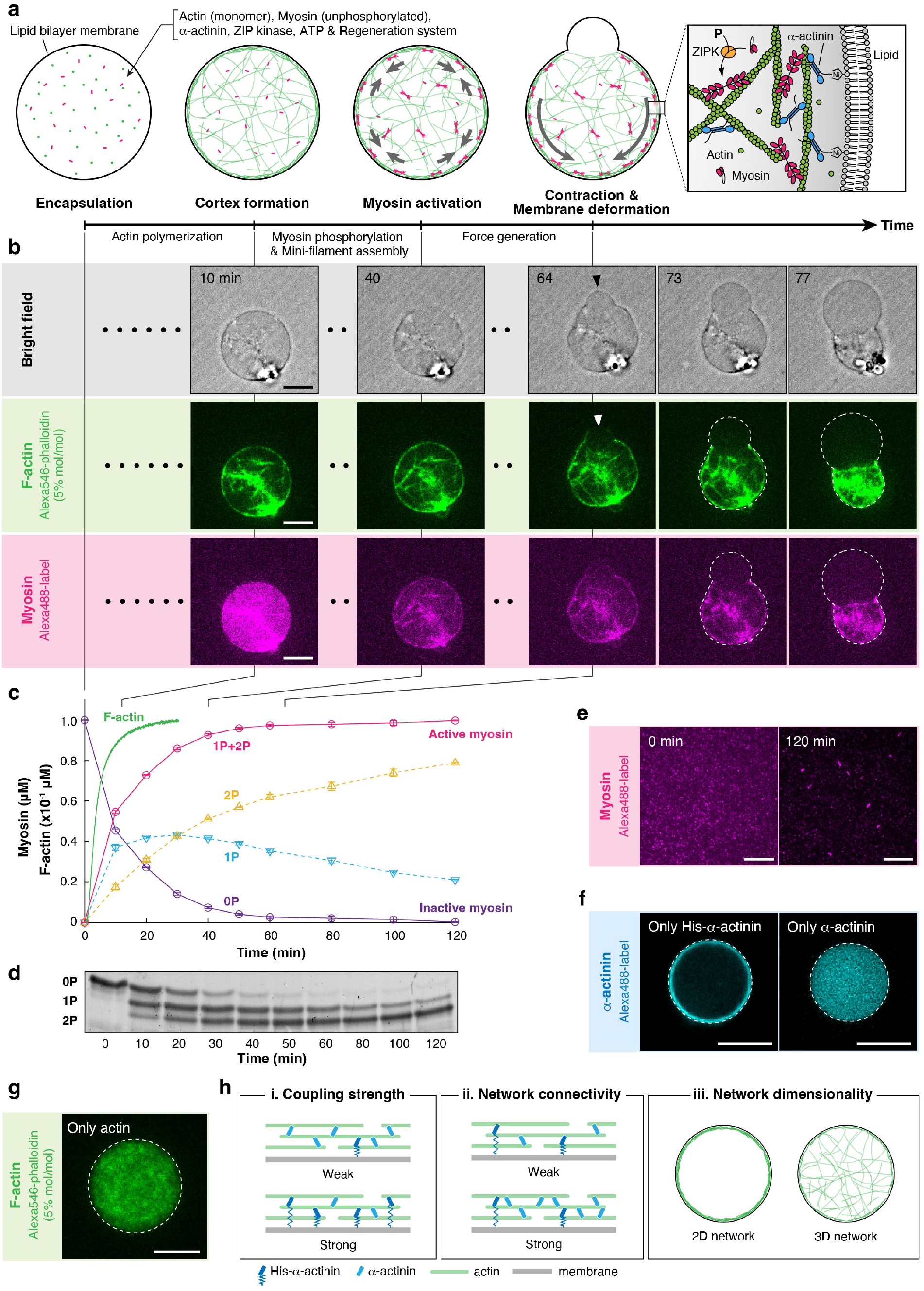
Reconstitution of the active actin cortex inside a cell-sized liposome. **a**, Schematic illustration of the experimental system. Actin monomers were mixed on ice with myosin, α-actinin (unphosphorylated), ZIP kinase, ATP, and ATP regeneration system. The mixture was immediately encapsulated into a cell-sized liposome by the inverted emulsion method, and the temperature was elevated to 25 ± 1°C to promote actin polymerization and activate myosin contractility. The membrane was composed of 85% phosphatidylcholine (PC), 10% phosphatidylglycerol (PG), and 5% Ni-NTA-conjugated lipid. **b**, Time-lapse confocal images of a liposome at the equator showing actin cortex formation, followed by myosin localization on F-actin, cortex contraction, and bleb formation and expansion (**Movie S1**). A black arrowhead and a white arrowhead indicate the bleb and the rupture point of the actin cortex, respectively. The liposome contains 10 μM actin, 2 μM His-α-actinin, 1 μM SMM (10% Alexa488-labeled), and 1.4 × 10^−4^ unit ml^-1^ ZIPK. **c**, Biochemical assays showing the kinetics of actin polymerization and myosin phosphorylation. Actin polymerization was monitored using pyrene-labeled actin. Myosin phosphorylation was quantified from gel electrophoresis shown in **d**. Three independent experiments were performed. Error bars indicate SDs. **d**, The image of urea/glycerol PAGE. 1 μM myosin was incubated with 1.4 × 10^−4^ unit ml^-1^ ZIPK in the presence of 1 mM ATP for the indicated times at 25°C. Unphosphorylated (0P), monophosphorylated (1P), and diphosphorylated (2P) myosin regulatory light chains were separated by gel electrophoresis. **e**, Total internal reflection fluorescence (TIRF) images of myosin before (left) and after (right) 120 min incubation with 1.4 × 10^−4^ unit ml^-1^ ZIPK in the presence of 1 mM ATP and ATP regeneration system. Submicrometer-long mini-filaments were formed by phosphorylation (**Fig. S1b**). **f**, Confocal images of liposomes at the equator containing only (left) his-tagged or (right) his-tag truncated α-actinin, showing that α-actinin was anchored to the membrane through the his-tag and Ni-NTA interaction, as illustrated in the inset of **a. g**, Confocal images of a liposome at the equator containing only actin, showing that F-actin has no specific interaction with the membrane. **h**, Schematic illustration of three physical parameters examined in the experiments. For all microscopic images, dashed lines indicate the liposome periphery. Scale bars, 10 μm.

F-actin in the cortex is inter-connected by various crosslinking proteins, including α-actinin, fimbrin, plastin, and fascin. The filaments are also anchored to the plasma membrane by diverse actin-membrane linkers such as ezrin-radixin-moesin (ERM) proteins and anillin^1^. Among the crosslinking proteins, α-actinin is the most abundant actin crosslinker in the actin cortex, as revealed by the proteomic analysis^36,37^, and is involved in diverse morphological transition processes, including cell motility^38,39,^ division^40^, and mitotic cell rounding^41^. We therefore used α-actinin to create a crosslinked network within a liposome. α-Actinin also binds to the plasma membrane through its pleckstrin-homology (PH), which has affinity for phosphatidylinositol 4,5-bisphosphate (PI(4,5)P_2_)^42,43^. To control this interaction, we instead used a defined linkage between a histidine-tag introduced to the recombinant α-actinin and Ni-NTA-conjugated phospholipids in the membrane (**Fig. 1a**, inset). We confirmed that, while his-tagged α-actinin (hereafter “His-α-actinin”) binds to the membrane (**Fig. 1f**, left), his-tag-truncated α-actinin (hereafter “α-actinin”) does not bind to the membrane (**Fig. 1f**, right). We also confirmed that F-actin shows no specific interactions with the membrane (**Fig. 1g**).

In subsequent experiments, we fixed the concentrations of actin (*C*_A_) and myosin (*C*_M_), and ZIPK at 10 μM, 1 μM, and 1.4 × 10^−4^ unit ml^-1^, respectively. Then, by varying the concentrations of His-α-actinin (*C*_H_) and α-actinin (*C*_N_), we independently controlled actin-membrane coupling strength, defined as *R*_C_ = *C*_H_/*C*_A_ (**Fig. 1h**, i) and actin network connectivity, defined as *R*_X_ = *C*_X_/*C*_A_ (**Fig. 1h**, ii), where *C*_X_ = *C*_H_ + *C*_N_ is the total crosslinker concentration. A co-sedimentation assay showed that the apparent dissociation constant *K*_d_ between His-α-actinin and F-actin is 1.4 μM, with binding to 10 μM actin reaching saturation at ∼5 μM (**Fig. S1c,d**). Therefore, we set the maximum *C*_X_ at 5 μM (*R*_X_ = 0.5). Furthermore, the spatial distribution of F-actin was modulated by adding methylcellulose to the inner solution of the liposomes. This crowding agent induces the localization of F-actin beneath the membrane by the depletion effect^44,45^, thereby enabling the formation of both a two-dimensional (2D) cortical network and a three-dimensional (3D) volume-spanning network (**Fig. 1h**, iii).

Taking all these advantages, we systematically investigated the effects of (i) actin-membrane coupling strength, (ii) actin network connectivity, and (iii) actin network dimensionality on the morphological transition process of cell-sized liposomes. The size of observed liposomes was 19.9 ± 7.56 μm (mean ± S.D.; *n* = 355) in diameter, comparable to the typical cell size.

### His-α-actinin concentration determines the morphological transition modes of liposomes

We initially focused on a 2D cortical network created with methylcellulose (**Fig. 2a**). Only His-α-actinin was used for simplicity (i.e., *R*_X_ = *R*_C_). First, we confirmed without myosin and ZIPK that His-α-actinin was co-localized with F-actin (**Fig. S2a**), and the actin density in a 2D cortical network remained nearly constant across the entire range of *C*_H_ (**Fig. 2b,c, Fig. S2b**). Then, we performed time-lapse microscopy in the presence of myosin and ZIPK to examine how varying *C*_H_ affects the morphological transition of liposomes (**Fig. 2d-g**). In the absence of His-α-actinin (*C*_H_ = 0; *R*_X_ = *R*_C_ = 0), neither global contraction of cortical actin networks nor detectable deformation of liposomes was observed (**Fig. 2e, Movie S2**). This result is consistent with previous reports, as myosin filaments are unable to induce global contraction of actin networks without actin crosslinking proteins at physiological concentration of ATP^46,47^,48 (maintained at 1 mM in our experiments). At *C*_H_ = 0.5 μM (*R*_X_ = *R*_C_ = 0.05), the cortical actin network showed contraction in 58% (14/24) of the liposomes, consistent with the lower limit α-actinin density (*R*_X_ = 0.05) required for the global contraction in millimeter-scale actomyosin gels^48^. More than half of these liposomes (64%) (9/14) showed detectable deformation during contraction (**Fig. 2f, Movie S3**). A further increase in *C*_H_ increased the fraction of liposomes showing detectable deformation from 38% (*C*_H_ = 0.5 μM; *R*_X_ = *R*_C_ = 0.05) to 100% (*C*_H_ = 5 μM; *R*_X_ = *R*_C_ = 0.5) (**Fig. 2g, Movie S4**).

**Figure 2.**
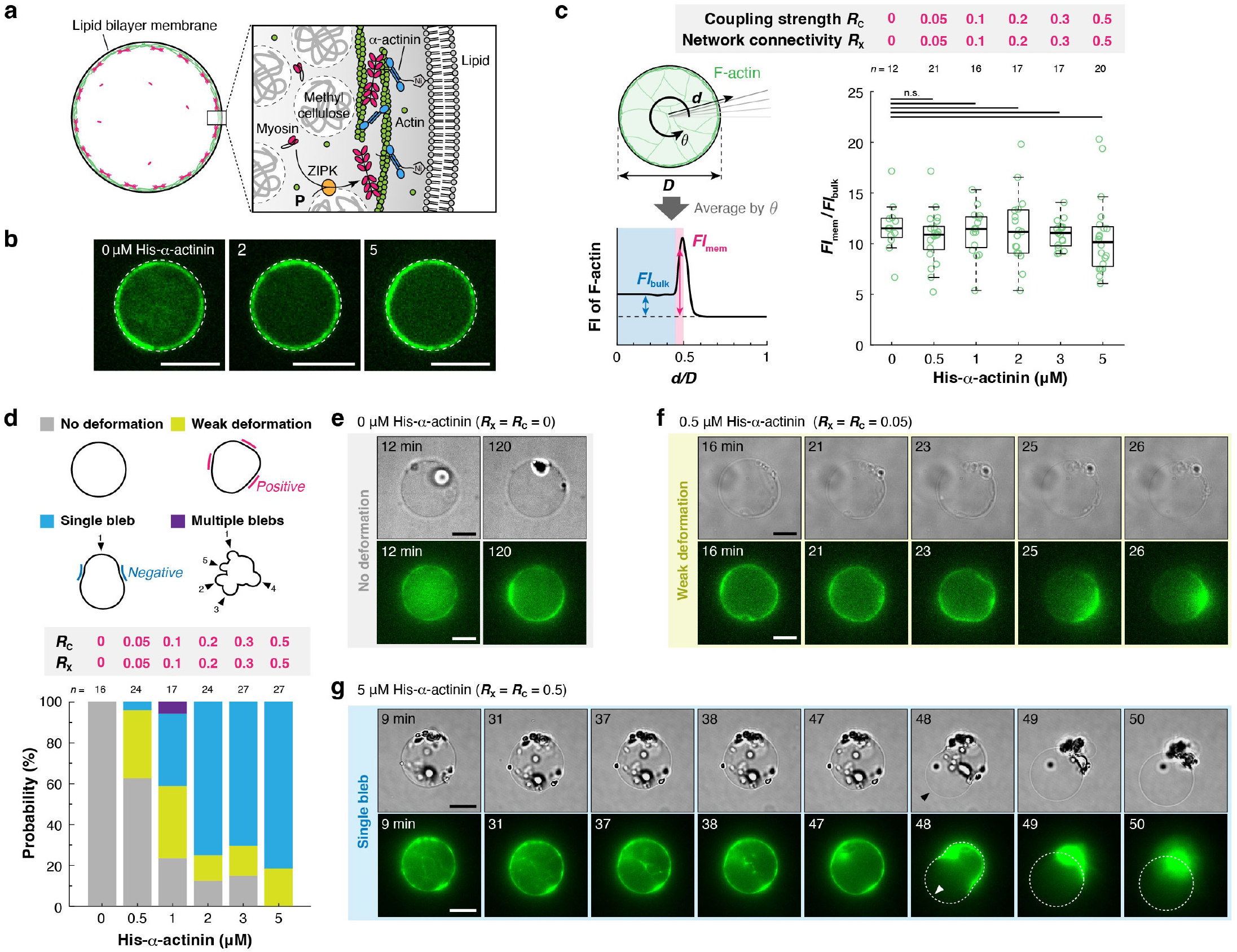
His-α-actinin concentration switches the morphological transition modes. **a**, Schematic illustration of the experimental system. Methylcellulose was added to localize F-actin beneath the membrane. **b**, Representative confocal images of liposomes at the equator showing F-actin distribution in the presence of various concentrations of His-α-actinin. **c**, Quantitative analysis of F-actin distribution. The fluorescence intensity of F-actin beneath the membrane *FI*_mem_ divided by the fluorescence intensity of F-actin in the bulk *FI*_bulk_ was plotted for each liposome. n.s.: *p* ≥0.05 (Welch’s *t*-test, two-sided). **d**, Probability distribution of the morphological transition modes of liposomes. Liposomes were classified into four categories in terms of their shapes: “No deformation”, “Weak deformation”, “Single bleb”, and “Multiple blebs”. Liposomes exhibiting negative membrane curvature at more than one location during the observation period were considered to have undergone bleb formation. These were further divided into “Single bleb” or “Multiple blebs” based on the number of blebs. Liposomes that showed detectable deformation without regions of negative curvature were classified as “Weak deformation”, while those showing no detectable deformation during the period of observation (2 hours) were assigned to “No deformation”. **e-g**, Time-lapse epifluorescence images of representative liposomes with various concentrations of His-α-actinin. **e**, No deformation (**Movie S2**), **f**, Weak deformation (**Movie S3**), and **g**, Single bleb (**Movie S4**). A black arrowhead and a white arrowhead indicate the bleb and the rupture point of the actin cortex, respectively. For all microscopic images, dashed lines indicate the liposome periphery. Scale bars, 10 μm.

To elucidate a relationship between *C*_H_ and the shape of liposomes, we classified the shape into four categories: “No deformation”, “Weak deformation”, “Single bleb”, and “Multiple blebs” (**Fig. 2d**, top). Quantitative analysis revealed that increasing *C*_H_ shifted the dominant mode from “No deformation” to “Weak deformation” to “Single bleb” (**Fig. 2d**, bottom). At the saturated *C*_H_ (5 μM; *R*_X_ = *R*_C_ = 0.5), all liposomes showed deformation, and 81% of liposomes (22/27) formed “Single bleb” (**Fig. 2d**, bottom). Formation of “Multiple blebs” was a very rare event; it was observed in only one case (1/17) at *C*_H_ = 1 μM (*R*_X_ = *R*_C_ = 0.1) (**Fig. S3a, Movie S5**).

### Actin-membrane coupling strength is a primary determinant controlling the morphological transition mode and the magnitude of membrane deformation

We demonstrated that the single parameter, *C*_H_, switched the morphological transition mode of liposomes (**Fig. 2**). However, changing *C*_H_ modulates both actin-membrane coupling strength (*R*_C_) and actin network connectivity (*R*_X_). Therefore, it remains unknown how these two physical parameters contribute to the morphological transition mode and magnitude of membrane deformation. To dissect the role of each parameter, we mixed His-α-actinin and α-actinin at various concentrations to control these two parameters independently (**Fig. 3a**). Then, we quantified the morphology and magnitude of membrane deformation in various conditions. We confirmed that the density of F-actin beneath the membrane was similar regardless of *C*_H_ and *C*_N_ (**Fig. 3b,c, Fig. S2b,c**). Therefore, this experimental setup enabled us to independently control the actin-membrane coupling strength and the network connectivity without a significant change in F-actin distribution.

**Figure 3.**
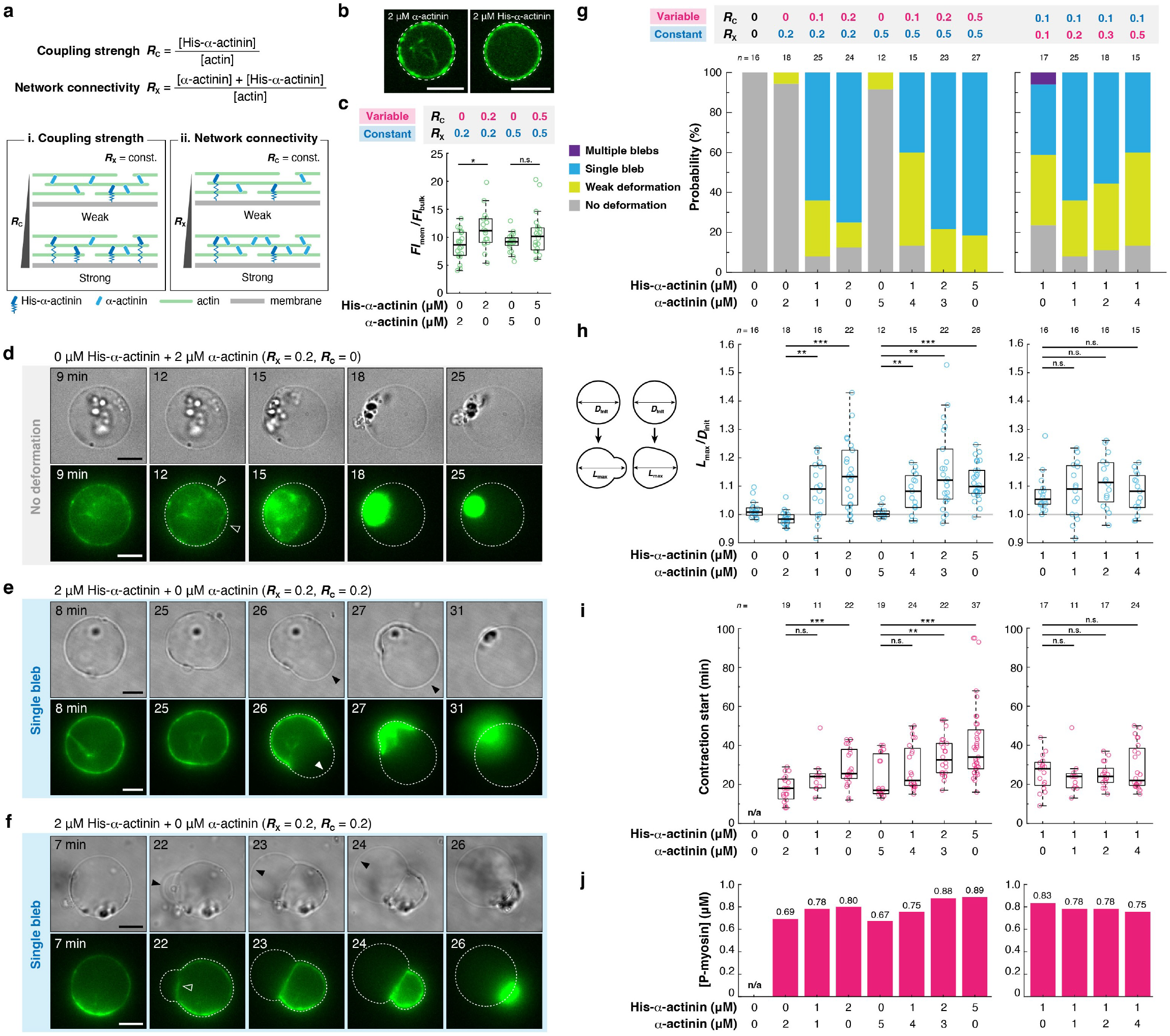
Actin-membrane coupling strength is the key parameter controlling the morphological transition mode and the magnitude of membrane deformation. **a**, Schematic illustration of the method to independently control actin-membrane coupling strength, *R*_C_, and network connectivity, *R*_X_. **b**, Representative confocal images of liposomes at the equator showing F-actin distribution in the presence of (left) α-actinin or (right) His-α-actinin. **c**, Quantitative analysis of F-actin distribution. The fluorescence intensity of F-actin beneath the membrane *FI*_mem_ divided by the fluorescence intensity of F-actin in the bulk *FI*_bulk_ was plotted for each liposome. **d-f**, Time-lapse epifluorescence images of representative liposomes under various conditions. **d**, The actin cortex was detached from the membrane (white open arrowheads). No clear membrane deformation was observed during the contraction (**Movie S6**). **e**, Rupture of the actin cortex (white filled arrowhead), followed by the contraction and blebbing of the membrane (black filled arrowheads) (**Movie S7**). **f**, Detachment of the actin cortex from the membrane (open white arrowhead), followed by the contraction and blebbing of the membrane (black filled arrowheads) (**Movie S8**). The cortex was not ruptured until the completion of contraction. **g-j**, Effects of *R*_C_ and *R*_X_ on (**g**) the morphological transition mode, (**h**) the magnitude of membrane deformation, (**i**) the contraction start time, and (**j**) the critical concentration of myosin required for the contraction, estimated from the median contraction start time in **i** and the time course of myosin phosphorylation (**Fig. 1c**). **g**, Liposomes were classified into four categories in terms of their shapes, as shown in **Fig. 2d. h**, The magnitude of membrane deformation is defined as the maximum length of the major axis of the liposome during contraction, *L*_max_, divided by the diameter of the liposome before contraction, *D*_init_, as illustrated on the left. For all microscopic images, dashed lines indicate the liposome periphery. Scale bars, 10 μm. ***: *p* < 0.001, **: *p* < 0.01, n.s.: *p* ≥0.05 (Welch’s *t*-test, two-sided).

Using this strategy, we first examined the effects of a variation in *R*_C_ while keeping *R*_X_ fixed (**Fig.3a**, i). In the absence of actin-membrane coupling (*R*_C_ = 0), most liposomes showed no detectable deformation, despite noticeable contraction of the cortical actin network at both *R*_X_ = 0.2 and 0.5 (**Fig. 3d, Fig. 3g**, left, **Movie S6**). In the presence of actin-membrane coupling (*R*_C_ > 0), liposomes were deformed by network contraction (**Fig. 3e,f, Fig. 3g**, left, **Fig. S3b, Movie S7, Movie S8, Movie S9**). With an increase in *R*_C_, the dominant morphological transition mode shifted from “No deformation” to “Weak deformation”, and then to “Single bleb” at both *R*_X_ = 0.2 and 0.5, which is consistent with the experiment using only His-α-actinin (**Fig. 2d**). At both *R*_X_ = 0.2 and 0.5, the magnitude of membrane deformation increased with higher *R*_C_ and reached a plateau at high *R*_C_-values (**Fig. 3h**, left). The onset time of network contraction was delayed with increasing *R*_C_ with both *R*_X_ = 0.2 and 0.5 (**Fig. 3i**, left), showing a trend similar to the magnitude of deformation (**Fig. 3h**, left). Using the time course of myosin phosphorylation (**Fig. 1c,d**), we estimated the concentration of phosphorylated myosin at the onset time of contraction (**Fig. 3j**, left). The result showed that higher myosin activity was required to initiate membrane deformation with increased actin-membrane coupling strength. Reducing the ZIPK concentration significantly delayed the onset time of contraction (**Fig.ES4a, Movie S10**), but the estimated concentration of phosphorylated myosin at the onset time and the magnitude of membrane deformation were comparable between the two concentrations, further validating the estimation method (**Fig. S4b-f**).

We next examined the effects of a variation in *R*_X_ while keeping *R*_C_ fixed (**Fig.3a**, ii). We found that, in contrast to a change in *R*_C_, changing *R*_X_ did not show a clear shift of the morphological transition modes (**Fig. 3g**, right). The magnitude of membrane deformation was not affected either by *R*_X_ (**Fig. 3h**, right). Notably, the onset time of network contraction was similar, regardless of *R*_X_ (**Fig. 3i**, right), implying that critical myosin activity required for the membrane deformation was not strongly affected by the network connectivity in this parameter range (**Fig. 3j**, right). Taken together, these results demonstrate that actin-membrane coupling strength, rather than network connectivity, is the primary determinant of both the morphological transition mode and the magnitude of membrane deformation.

### Bleb formation is initiated by one of two distinct mechanisms, “Detachment” or “Rupture”, depending on the balance between actin-membrane coupling strength and network connectivity

Time-lapse observation revealed that bleb formation was initiated by two distinct mechanisms: “Detachment” or “Rupture” (**Fig. 4a**). As for the “Rupture,” bleb formation was initiated by the spontaneous rupture of the actin cortex beneath the membrane (**Fig. 3e**, filled arrowhead, **Movie S7**). By contrast, as for the “Detachment,” bleb formation was initiated by the local detachment of the actin cortex from the membrane (**Fig. 3f**, open arrowhead, **Movie S8**). Note that, in some cases showing “Detachment,” the cortex was eventually ruptured during the bleb expansion (**Fig. S3b**, filled arrowhead, **Movie S9**). We found that, at constant *R*_X_, “Detachment” was dominant at low *R*_C_, whereas “Rupture” became dominant with increasing *R*_C_ (**Fig. 4b**, left). On the contrary, at constant *R*_C_, “Rupture” was dominant at low *R*_X_, whereas “Rupture” became dominant with increasing *R*_X_ (**Fig. 4b**, right). These results suggest that the bleb initiation mechanism is determined by how well filaments are connected in the network and how strongly the network is coupled to the membrane.

**Figure 4.**
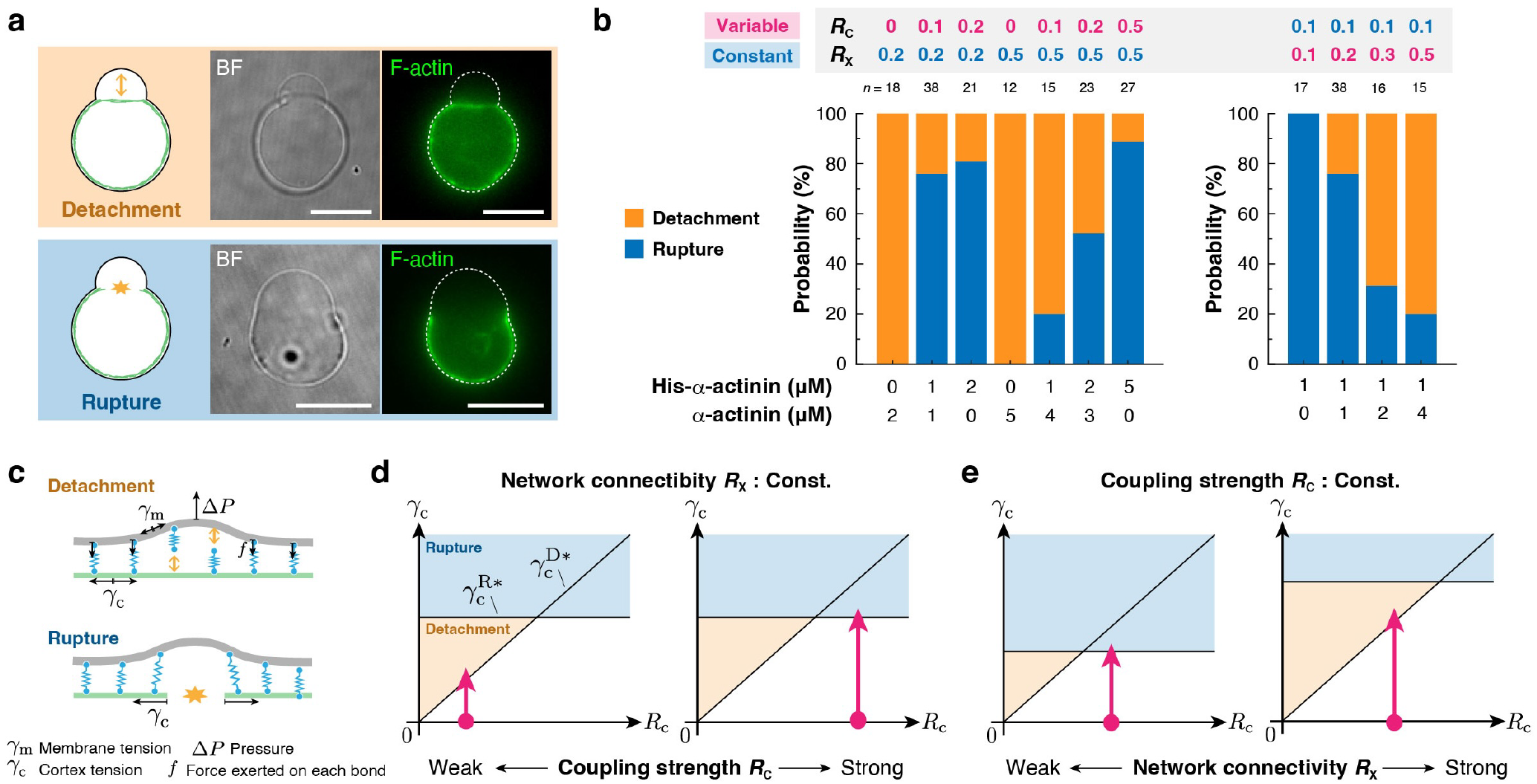
Bleb formation is induced by one of two distinct mechanisms, which is determined by the balance between actin-membrane coupling strength and network connectivity. **a**, Two types of bleb initiation mechanisms observed in the experiments. Top: A bleb is induced by the detachment of the cortical actin network from the membrane (**Movie S8, Movie S9**). Bottom: A bleb is induced by the rupture of the cortical actin network (**Movie S7**). Dashed lines indicate the liposome periphery. Scale bars, 10 μm. **b**, Probability distribution of the two types of bleb initiation mechanisms under various conditions. **c**, Schematic illustration of the molecular mechanisms underlying the two bleb initiation mechanisms. **d,e**, Phase diagrams of the bleb initiation mechanisms. 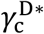 and 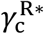 indicate the critical cortex tensions required for the detachment and rupture mechanisms, respectively. 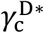 is proportional to the actin-membrane coupling strength *R*_C_, as shown in **Eq. 8**. By contrast, 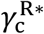 is an increasing function of the network connectivity *R*_X_, but it is independent of *R*_C_. **d**, Dependence on actin-membrane coupling strength, *R*_C_. **e**, Dependence on network connectivity, *R*_X_. Magenta arrows indicate the pathways taken by the system upon myosin phosphorylation.

To gain physical insight into the bleb initiation mechanism, we developed a simple theoretical model. First, we introduce a theoretical description of the “Detachment” mechanism (**Fig. 4c**, top). We assume that a hydrostatic pressure difference between the inside and outside of the liposome, Δ*P*, is balanced by the elastic restoring force, *F*, generated by the coupling between the actin cortex and the membrane-bound His-α-actinin molecules until the time point just before bleb initiation:

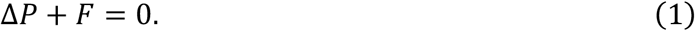

Using the Young-Laplace equation, Δ*P* can be written as

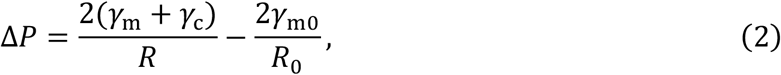

where *γ*_m_, *γ*_c_, and *R* indicate the membrane tension, the actin cortex tension, and radius of the liposome at a certain time point *t*, respectively. *γ*_m0_ and *R*_0_ indicate the membrane tension and the radius of the liposome right after the liposome formation (*t* = 0), respectively^14,20^. As myosin phosphorylation proceeds and the internal pressure reaches a level close to the bleb initiation threshold, the contribution of *γ*_m_ would become much smaller than that of *γ*_c_ (*γ*_c_ ≫ *γ*_m_), which was confirmed by *in silico* simulation (see **Fig. 6f, Fig. 7c,g,k**). Thus, the internal pressure can be approximated as

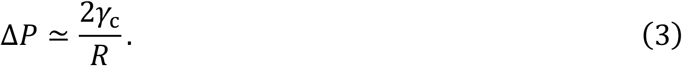

Next, we consider the binding and unbinding dynamics of actin-membrane coupling. Although there are two potential molecular pathways for decoupling – either unbinding of His-α-actinin from F-actin or unbinding of the His-tag from Ni-NTA-lipid – we assume that the former is the predominant pathway for simplicity (see **Supplementary Note 2**, Section 7). In the absence of the hydrostatic pressure (Δ*P* = 0), the actin-bound His-α-actinin density *ρ* is governed by the kinetic equation:

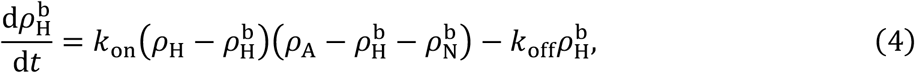

where 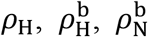, and *ρ*_A_ indicate the surface densities of His-α-actinin, His-α-actinin bound on F-actin, α-actinin bound on F-actin, and F-actin beneath the membrane, respectively. *k*_on_ and *k*_off_ are the rate constants for binding and unbinding between His-α-actinin and F-actin, respectively. Considering that Δ*P* tends to separate the membrane from the actin cortex, and assuming that each His-α-actinin-F-actin bond is equally stressed, the force per bond, *f*, becomes 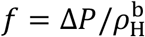. Using Bell’s kinetics model^49^, the kinetic equation (**Eq. 4**) is replaced by

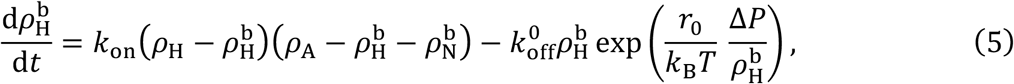

where 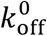 is the zero-force unbinding rate constant, and *r*_0_ is a characteristic distance to overcome the energetic barrier for bond separation^49,50,51^. When a small constant Δ*P* is applied, the unbinding rate (second term on the right-hand side of **Eq. 5**) transiently increases, leading a slight decrease in 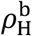, and then the system reaches a new equilibrium 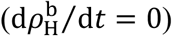. In contrast, when a large Δ*P* is applied, the unbinding rate (second term) becomes much larger than the binding rate (first term), causing 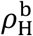 to rapidly approach zero. There will thus exist the critical pressure, Δ*P*^*^, that is just sufficient to detach all His-α-actinin molecules from F-actin within a local region (**Supplementary Note 1**). If it is assumed that F-actin near the membrane retains many free binding sites for α-actinin 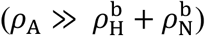, we obtain the critical pressure:

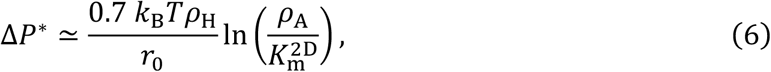

where 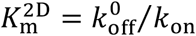 is the dissociation constant of His-α-actinin from F-actin in a 2D network (**Supplementary Note 1**). Therefore, using **Eq. 3**, the critical cortical tension required for the “Detachment” mechanism is estimated as

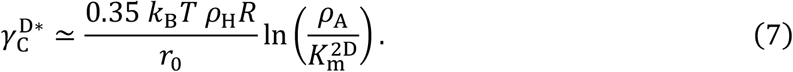

Since *R*_C_ ≃ *ρ*_A_/*ρ*_A_, this tension is proportional to the actin-membrane coupling strength *R*_C_:

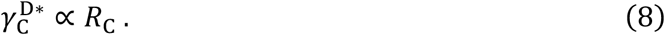

We next consider the “Rupture” mechanism (**Fig. 4c**, bottom). It can be reasonably approximated that the critical cortical tension required for the actin network rupture 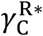 is an increasing function of *R*_X_ and independent of *R*_C_ in our parameter range^52^.

Based on the analytical results, we generated phase diagrams predicting which bleb initiation mechanism is selected (**Fig. 4d,e**). At constant *R*_X_, the model predicts that the mechanism shifts from “Detachment” to “Rupture” with increasing *R*_C_ (**Fig. 4d**). At constant *R*_C_, the model predicts that the mechanism shifts from “Rupture” to “Detachment” with increasing *R*_X_ (**Fig. 4e**). These predictions are entirely consistent with the experimental results (**Fig. 4b**). Furthermore, the model predicts that at constant *R*_X_, higher myosin activity is required to initiate bleb formation at higher *R*_C_. This is also consistent with the experimental results (**Fig. 3i**, left, **Fig. 3j**, left). Collectively, this simple theoretical model explains that the mechanism triggering the bleb initiation is regulated by the balance between actin-membrane coupling strength and network connectivity.

### 3D volume-spanning actin network facilitates the formation of multiple blebs

In cells, F-actin not only forms a dense cortex beneath the plasma membrane but also forms a sparse volume-spanning network throughout the cytoplasm. Therefore, it is important to assess how this coexisting bulk network influences membrane deformation. To address this, we created a volume-spanning 3D network by excluding methylcellulose from the inner solution (**Fig. 5a**) and examined how the concentration of His-α-actinin (*C*_H_) affects morphological transitions, as analyzed in the 2D network (**Fig. 2**). As in the 2D network case, only His-α-actinin was used for simplicity, (i.e., *R*_X_ = *R*_C_). We first confirmed that an increase in *C*_H_ increased the cortical F-actin density, reaching a plateau at *C*_H_ = 2 μM (*R*_X_ = *R*_C_ = 0.2) (**Fig. 5b,c, Fig. S2d**). However, the relative cortical F-actin density *FI*_mem_/*FI*_bulk_ was lower than that in the cases with methylcellulose (**Fig.2c, Fig. 3c, Fig. S2e**), suggesting that F-actin formed a sparse network in the bulk region even at high *C*_H_. We next performed time-lapse microscopy to examine how *C*_H_ affects the morphological transition of liposomes (**Fig. 5d-h**). In the absence of His-α-actinin (*C*_H_ = 0 μM; *R*_X_ = *R*_C_ = 0), neither contraction of actin networks nor deformation of liposomes was observed (**Fig. 5e, Movie S11**). At *C*_H_ = 0.5 μM (*R*_X_ = *R*_C_ = 0.05), the actin network showed contraction in 39% (7/18) of the liposomes, and some of these liposomes deformed during the contraction (**Fig. 5f, Movie S12**). At *C*_H_ = 1 μM (*R*_X_ = *R*_C_ = 0.1), the actin network showed contraction in all the liposomes (20/20). A further increase in *C*_H_ induced larger deformation (**Fig. 5g,h Movie S13, Movie S14**).

**Figure 5.**
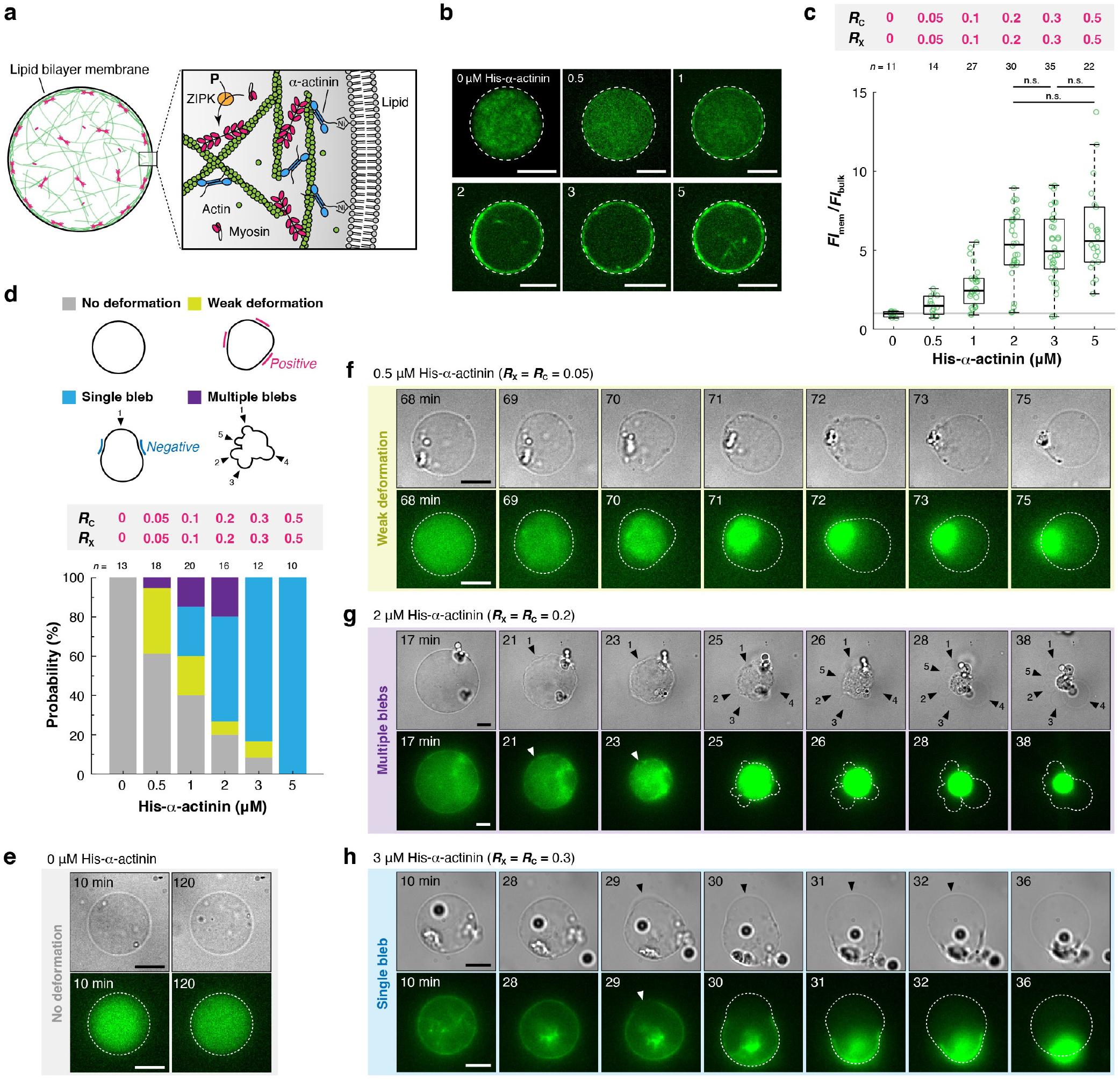
3D volume-spanning actin network facilitates the formation of multiple blebs. **a**, Schematic illustration of the experimental system. Methylcellulose was excluded from the inner solution to create a 3D volume-spanning actin network. **b**, Representative confocal images of liposomes at the equator showing F-actin distribution in the presence of various concentrations of His-α-actinin. **c**, Quantitative analysis of F-actin distribution. The fluorescence intensity of F-actin beneath the membrane *FI*_mem_ divided by the fluorescence intensity of F-actin in the bulk *FI*_bulk_ was plotted for each liposome. n.s.: *p* ≥ 0.05 (Welch’s *t*-test, two-sided). **d**, Probability distribution of the morphological transition modes of liposomes. Liposomes were classified into four categories in terms of their shapes, as in the cases with a 2D cortical network (**Fig. 2d, Fig. 3g**). **e-h**, Time-lapse epifluorescence images of representative liposomes under various concentrations of His-α-actinin. **e**, No deformation (**Movie S11**), **f**, Weak deformation (**Movie S12**), **g**, Multiple blebs (**Movie S13**), and **h**, Single bleb (**Movie S14**). Black arrowheads and white arrowheads indicate the blebs and the rupture points of the actin cortex, respectively. For all microscopic images, dashed lines indicate the liposome periphery. Scale bars, 10 μm.

**Figure 6.**
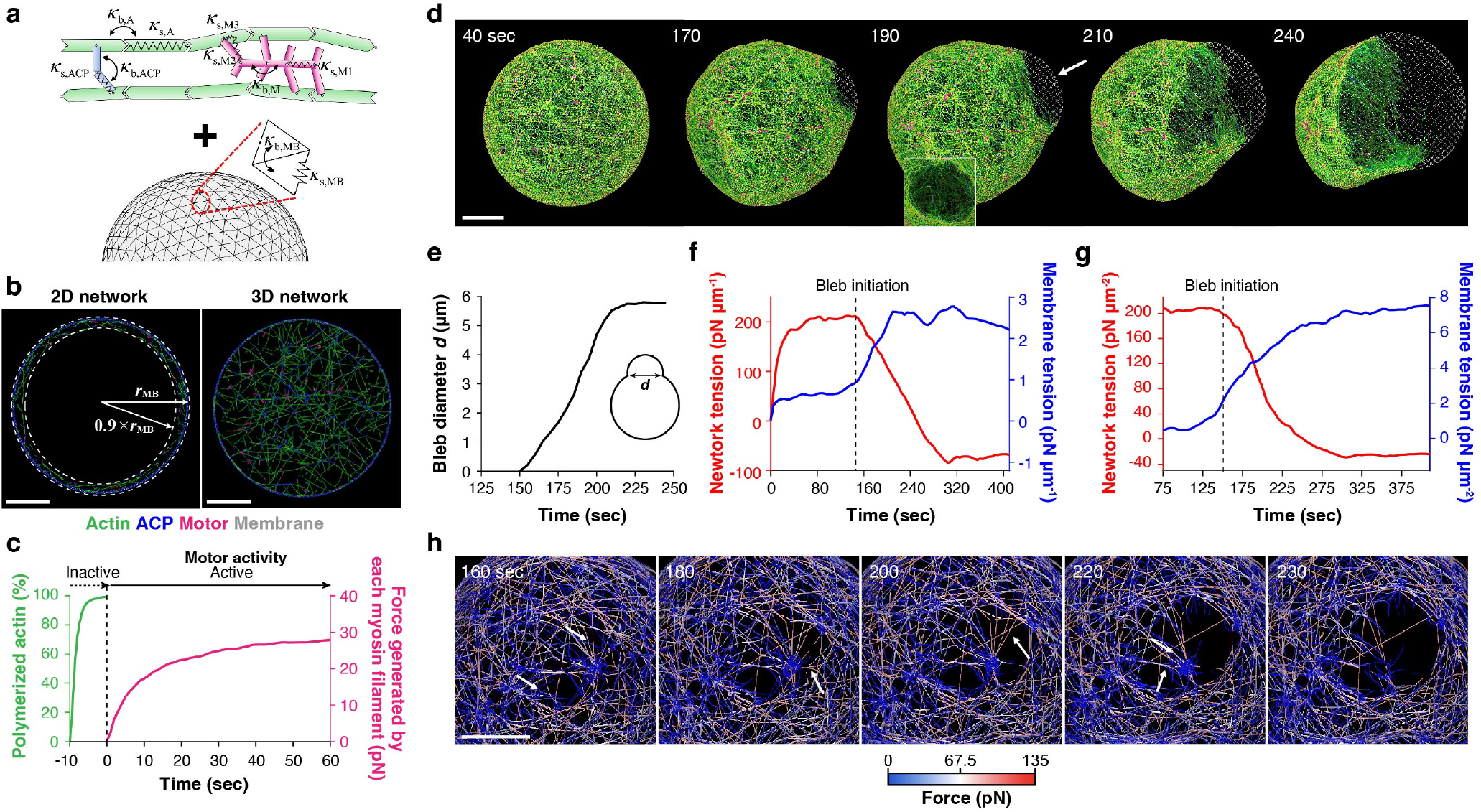
Agent-based computational model for dissecting the molecular mechanism of bleb formation and tension dynamics originating from the dynamic interplay of an actin network and a membrane. **a**, The model comprises an actin network encapsulated by a deformable membrane. **b**, Cross-sectional views of (left) 2D and (right) 3D networks assembled within the membrane. For the 2D network, the network was assembled near the membrane within a space between *r* = 0.9 × *r*_MB_ and *r* = *r*_MB_, where *r*_MB_ is a radial distance from the initial membrane center. The mean length of F-actin in the 2D network and 3D network was ∼4.2 μm and ∼5.2 μm, respectively. **c**, Time courses of actin polymerization and motor activity. These were estimated by measuring F-actin concentration and force generated by each myosin filament, respectively. Myosin motors were activated after network formation in an all-or-none fashion to reduce computational cost. This likely has minimal effects on network contraction and membrane deformation, as significant time was required to develop network tension before bleb initiation, as shown in **f. d**, Example of single bleb formation observed under the reference condition (*R*_X_ = *R*_C_ = 0.08, motor density *R*_M_ = 0.01) (**Movie S16**). The bleb started emerging from ∼150 s in this example, followed by expansion. The inset shown at 190 s represents a hole on the network beneath the bleb in a different view as indicated by the arrow, showing that this bleb was initiated by a network rupture. **e**, The diameter of the bleb measured over time. **f**, Global tension acting on the network (red) and the membrane (blue) over time. **g**, Local network and membrane tension measured near the bleb over time. **h**, Time-lapse images showing that catastrophic severing events of F-actins indicated by arrows led to the initial formation and expansion of a rupture on the network (**Movie S17**). Scale bars, 2 μm.

**Figure 7.**
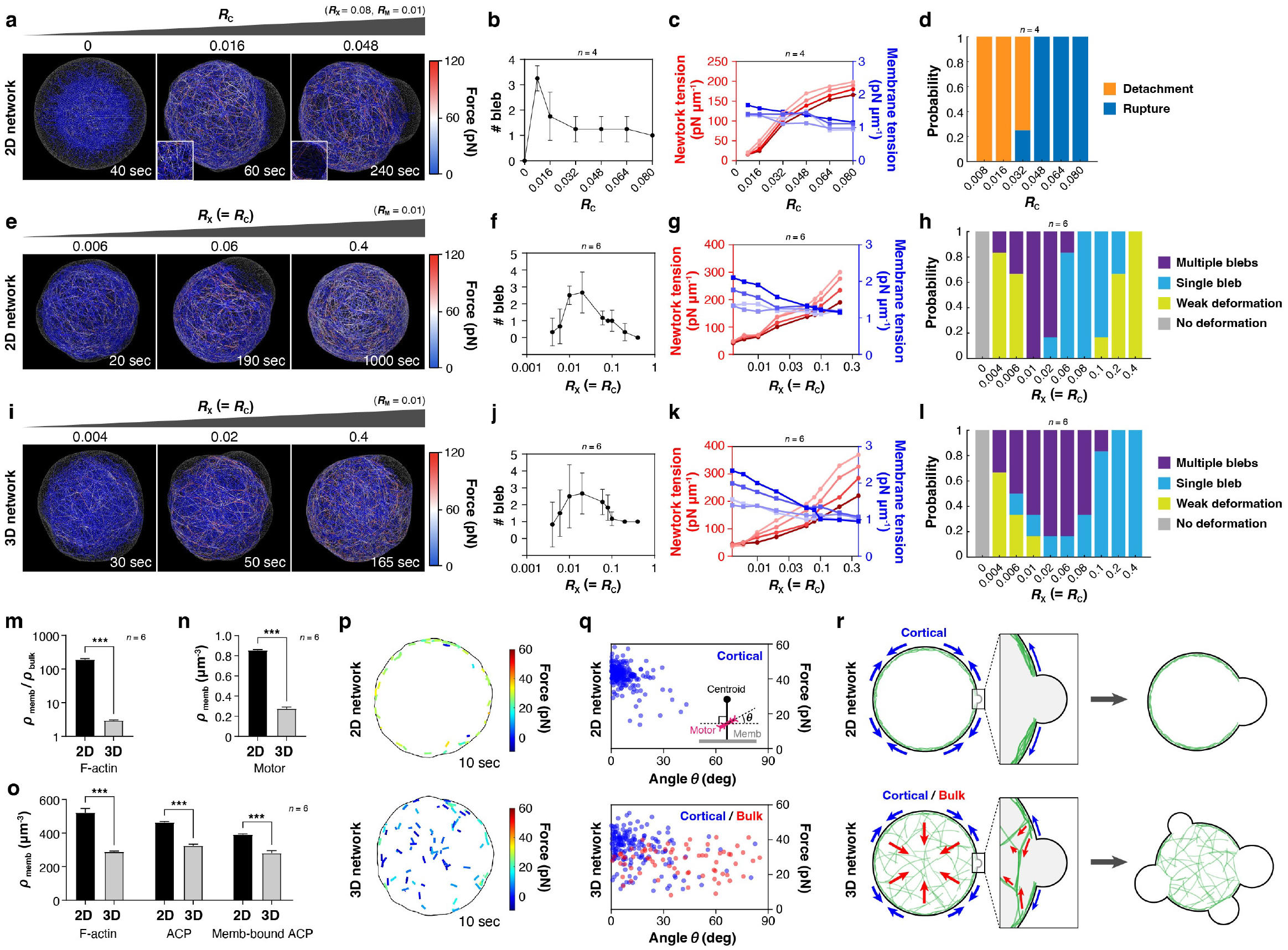
Different patterns of shape changes and bleb formation emerged, depending on the actin-membrane coupling strength, network connectivity, and the network dimensionality. **a-d**, Effects of actin-membrane coupling strength, *R*_C_ with the 2D network. *R*_C_ was changed with *R*_X_ fixed at the reference value (0.08). **e-h**, Effects of membrane-bindable ACP density (i.e., *R*_X_ = *R*_C_) with the 2D network. **i-l**, Effects of membrane-bindable ACP density with the 3D network. **a,e,i**, Representative snapshots of the network and membrane under different conditions (**Movies S19-S24**). Color indicates force magnitude on each F-actin. (**a**, left) Without actin-membrane coupling (*R*_C_ = 0), the network contracted into a small cluster without noticeable membrane deformation. (**a**, center) At *R*_C_ = 0.016, bleb formation was initiated by detachment of the network from the membrane, evidenced by the intact network remaining beneath the bleb (inset). (**a**, right) At *R*_C_ = 0.048, bleb formation was initiated by network rupture, evidenced by a hole in the network beneath the bleb (inset). **b,f,j**, Number of blebs as a function of (**b**) *R*_C_ and (**f,j**) *R*_X_ (= *R*_C_). Error bars indicate SDs. **c,g,k**, Global network/membrane tension before and after bleb formation as a function of (**c**) *R*_C_ and (**g,k**) *R*_X_ (= *R*_C_). Darker colors show tension at later times. **d**, Effect of *R*_C_ on the bleb initiation mechanism. **h,l**, Effect of membrane-bindable ACP density (*R*_X_ = *R*_C_) on morphological transition modes, with the (**h**) 2D or (**l**) 3D networks. **m**, Ratio of F-actin density near the membrane (*ρ*_memb_) to F-actin density in the bulk (*ρ*_bulk_) in the 2D and 3D networks under the reference condition (*R*_X_ = *R*_C_ = 0.08, *R*_M_ = 0.01). **n**, Density of motors beneath the membrane in the 2D and 3D networks (*R*_X_ = *R*_C_ = 0.08, *R*_M_ = 0.01). **o**, Densities of F-actin, ACP, and membrane-bound ACPs beneath the membrane in the 2D and 3D networks (*R*_X_ = *R*_C_ = 0.08, *R*_M_ = 0.01). **p,q**, Orientation of myosin filaments and the magnitude of motor forces in the 2D network (**Movie S25**) and the 3D network (*R*_X_ = *R*_C_ = 0.08, *R*_M_ = 0.01) (**Movie S26**) (**p**) visualized at a single time point and (**q**) the quantitative analysis results. **r**, Schematic representation of the force distribution in the 2D and 3D networks. Arrows indicate force directions, and the lengths represent magnitudes.

Similar to the 2D network (**Fig. 2d**), an increase in *C*_H_ shifted the dominant mode from “No deformation” to “Weak deformation” to “Single bleb” (**Fig. 5d**). Indeed, at the saturated *C*_H_ (5 μM; *R*_X_ = *R*_C_ = 0.5), all liposomes (10/10) formed single blebs (**Fig. 5h, Movie S14**). Remarkably, at intermediate *C*_H_ (0.5, 1, and 2 μM; *R*_X_ = *R*_C_ = 0.05, 0.1, and 0.2), some liposomes exhibited “Multiple blebs” (**Fig. 5g, Movie S13**), reminiscent of the morphological transition observed during apoptosis^5^. This is a distinct feature from the 2D network conditions (**Fig. 2d, Fig. 3g**).

Furthermore, as demonstrated in **Fig. 3** and **Fig. 4**, we systematically investigated the individual contributions of *R*_C_ and *R*_X_ in the 3D network by varying *C*_H_ and *C*_N_ independently (**Fig. S5**). We confirmed that the basic two features revealed under the 2D network conditions (**Fig. 3, Fig. 4**) are preserved in the 3D network conditions: increasing *R*_C_ shifted the dominant morphological transition mode from “No deformation” to “Weak deformation” and then to “Single bleb” (**Fig. S5a,c**), and increasing *R*_X_ shifted the dominant bleb initiation mechanism from “Rupture” to “Detachment” (**Fig. S5b,d**). These parametric explorations further corroborate the universality of the regulatory mechanisms identified in the minimal reconstituted system. Moreover, we revealed that the 3D network with intermediate *R*_C_ and *R*_X_ is the optimal condition for multiple bleb formation (**Fig. S5a,c**).

### Catastrophic F-actin severing initiates bleb formation by the “Rupture” mechanism

Although our theoretical model could predict which mechanism is likely to trigger bleb initiation depending on the two physical parameters, how bleb initiation occurs at a molecular scale remains elusive. To define the molecular mechanism, we developed an agent-based model with an actin network encapsulated by a deformable membrane (**Fig. 6a, Supplementary Note 2**, Sections 1-8). The network is composed of F-actin, myosin mini-filaments, and actin crosslinking proteins (ACPs) with the geometry analogous to α-actinin. A fraction of ACPs can reversibly bind to the membrane as His-α-actinin used in experiments. The actin concentration was 10 μM in all simulations to be consistent with experiments. Unless specified, the membrane diameter was 8 μm, smaller than the mean diameter used in experiments (∼20 μm) to reduce the computational cost. We confirmed that the results obtained using a system with a diameter of 16 μm were similar (**Fig. S6, Fig. S7, Movie S15**). As in the experiments, we created both a 2D cortical network (**Fig. 6b**, left) and a volume-spanning 3D network (**Fig. 6b**, right) and then compared the morphological transition process.

First, we performed simulations using the 2D network (**Fig. 6b**, left). Initially, actin nucleation and polymerization took place beneath the membrane in the presence of ACPs and myosin filaments to let them self-organize into the 2D network (**Fig. 6c**, green line). After the cortex formation, myosin was activated to start generating forces (**Fig. 6c**, magenta line). In the reference case with *R*_X_ = *R*_C_ = 0.08, which is close to 1 μM His-α-actinin condition (*R*_X_ = *R*_C_ = 0.1) in the experiments (**Fig. 2**), a single bleb started forming at ∼150 s and then expanded substantially over time (**Fig. 6d,e, Fig. S6a-d, Movie S16**) as observed in the experiments. This bleb was initiated by a network rupture triggered by successive F-actin severing events induced by large tensile forces (**Fig. 6h, Fig. S8a, Movie S17**). Such rupture-initiated bleb formation was not observed when we did not incorporate F-actin severing in the model, indicating that severing is the essential process for the network rupture^53^ (**Supplementary Note 2**, Section 5).

The computational model enables us to measure the network tension and membrane tension separately. Global network tension increased and remained at a plateau (∼200 pN μm^-1^) for ∼70 s, after which it began to decrease upon bleb formation and expansion (**Fig. 6f**, red line, **Fig. S6e,f**). This plateau phase implies that it took significant time for large tensile forces to accumulate in a few filaments and reach the critical severing threshold^54^ (assumed to be 300 pN; **Supplementary Note 2**, Section 4). In contrast, global membrane tension was initially low and then increased during bleb expansion (**Fig. 6f**, blue line), as the expanding bleb membrane experienced expansile forces resulting from volume conservation. Local network/membrane tension measured near the bleb showed similar tendencies before and after bleb formation (**Fig. 6g**). The local network tension and local motor density were higher in the region showing bleb formation than those in the region showing only membrane deformation (**Fig. S9a,b**), implying that locally higher contractility induced by more motors enhanced network tension and thus led to the network rupture. This was verified by examining the impacts of motor density; higher motor density induced more frequent F-actin severing events and more blebs (**Fig. S10a-c**). The network rupture was also accompanied by a decrease in local membrane-bound ACP density (**Fig. S6g, Fig. S9c-e**). Taken together, the computational simulation revealed that catastrophic F-actin severing events at a local region are required for initiating bleb formation via the network rupture mechanism, and this is triggered by higher local motor density, followed by a local decrease in membrane-bound ACPs.

### Bleb is initiated by the “Detachment” mechanism when actin-membrane coupling is weak

Using this 2D network model, we next examined the effect of the actin-membrane coupling *R*_C_ by reducing it from the reference case, with the network connectivity *R*_X_ fixed at 0.08 (**Fig. 7a-d, Fig. S10d**). Without network-membrane coupling (*R*_C_ = 0), the network contracted into a smaller cluster without noticeable membrane deformation (**Fig. 7a**, left, **Fig. S10d**, left, **Movie S18**). At low network-membrane coupling (*R*_C_ = 0.008 or 0.016), a bleb was initiated by the detachment of the network from the membrane (**Fig. 7a**, center, **Fig. S10d**, center, **Movie S19**). With *R*_C_ higher than 0.048, bleb formation was driven by the network rupture as in the reference case (**Fig. 7a**, right, **Fig. S10d**, right, **Movie S20**). At the intermediate regime (*R*_C_ = 0.032), some blebs were induced by “Detachment”, but the rest was induced by “Rupture” (**Fig. 7d**). These results are consistent with the experiments (**Fig. 4b**, left). We found that the network tension increased nearly proportional to *R*_C_ up to the intermediate regime (*R*_C_ = 0.048) (**Fig. 7c**, red lines), whereas membrane tension remained constant regardless of *R*_C_ (**Fig. 7c**, blue lines). At higher *R*_C_, robust coupling between the membrane and network enhanced the network stability. Thereby, a greater force was required to rupture the network and initiate bleb formation, consistent with the experiments (**Fig. 3i**, left, **Fig. 3j**, left). At lower *R*_C_, the network began detaching from the membrane even before tension increased to a sufficiently high level required for F-actin severing (**Fig. 7c**, red lines, *R*_C_ ≤ 0.016), resulting in bleb formation by the detachment mechanism, consistent with the model prediction (**Fig. 4d**, left).

### Comparison between 2D and 3D networks reveals an active role of the bulk actin network

Our experiments showed that the 3D network induces the formation of multiple blebs more frequently (**Fig. 2d, Fig. 5d, Fig. S5**), implying the potential contribution of the sparse bulk actin network to the formation of multiple blebs. To dissect the underlying molecular mechanism, we ran simulations using both 2D and 3D networks (**Fig. 6b**) with various ACP densities and compared the outcomes (**Fig. 7e-l**). As in the case of experiments (**Fig. 2, Fig. 5**), all ACPs were allowed to bind to the membrane (*R*_X_ = *R*_C_), so changes in *R*_X_ also changed the density of actin-membrane coupling points.

With the 2D network in the absence of ACPs (*R*_X_ = *R*_C_ = 0), neither the network contraction nor membrane deformation was observed (**Fig. 7h**). At low ACP density (*R*_X_ = *R*_C_ = 0.006), either multiple small blebs or only membrane deformation without any bleb were observed (**Fig. 7e**, left, **Fig. 7f,h, Fig. S10e**, left, **Movie S21**). At intermediate ACP density (*R*_X_ = *R*_C_ = 0.06), only a single bleb appeared more likely than multiple blebs (**Fig. 7e**, center, **Fig. 7f,h, Fig. S10e**, center, **Movie S22**). At high ACP density (*R*_X_ = *R*_C_ = 0.4), blebs were negligibly small, which is attributed to high network connectivity (**Fig. 7e**, right, **Fig. 7f,h, Fig. S10e**, right, **Movie S23**); although small network ruptures followed by the emergence of tiny membrane bulges smaller than 0.8 μm in diameter were observed, they did not expand enough to initiate bleb formation because the ruptures did not increase in size due to a large number of cross-linking points between F-actins (**Fig. S8b**). While the initial membrane tension was rather independent of ACP density (**Fig. 7g**, blue lines), network tension increased nearly proportionally with ACP density, as more ACPs rendered the network more elastic by increasing both cross-linking points within the network and coupling points to the membrane (**Fig. 7g**, red lines).

Results obtained with the 3D network (**Fig. 6b**, right, **Fig. 7i-l, Fig. S10f**) were qualitatively similar to those obtained using the 2D network (**Fig. 7e-h, Fig. S10e**). The minimal ACP density required for the bleb formation was comparable between 2D and 3D networks (*R*_X_ = *R*_C_ = 0.004), whereas the maximum ACP density permitting bleb formation was higher in the 3D network (**Fig. 7h,l**). In the 3D network, membrane-bindable ACPs were distributed across the entire volume within the membrane, resulting in fewer effective coupling points between the network and the membrane. Consequently, the bleb formation probability was higher in the 3D network at the same *R*_X_ (= *R*_C_). We compared the densities of F-actin, myosin motors, and membrane-bound ACPs between the 2D and 3D networks for our reference case (*R*_X_ = *R*_C_ = 0.08), and confirmed that all of these components were present at lower densities beneath the membrane in the 3D network (**Fig. 7m-o**), consistent with the F-actin distribution observed in the experiments (**Fig. 2c, Fig. 5c, Fig. S2e**).

Importantly, with the 3D network, multiple blebs formed over a wider range of ACP densities (**Fig. 7h,l**, purple bars and **Movie S24**), reproducing the trend observed in the experiments (**Fig. 2d, Fig. 4d**). We found that the orientation angle of individual myosin filaments showed a marked difference between 2D and 3D networks, and not only the cortical myosin filaments but also myosin filaments in the bulk region generated significant forces (**Fig. 7p,q, Movie S25, Movie S26**). While myosin filaments beneath the membrane were oriented parallel to the membrane, which may contribute to the cortical tension development both in 2D and 3D networks (**Fig. 7q**, blue plots), myosin filaments in the bulk region of the 3D network showed nearly random orientations (**Fig. 7q**, red plots), which can generate inward pulling force to the membrane as an ensemble (**Fig. 7r**, left). At the molecular scale, sparse bundled actin networks in the bulk region generate contractile forces that act as antagonistic forces opposing membrane expansion (**Fig. 7r**, insets). This will trigger the formation of new blebs at different positions to release the elevated pressure in liposomes, resulting in multiple bleb formation more likely (**Fig. 7r**, right). Overall, although the cortical network is expected to play a central role in bleb formation, our results indicate that the bulk actin network may also play an important role in regulating the number of blebs in cells.

### Local perturbation on actin-membrane coupling strength or network connectivity controls bleb position

Previous studies in living cells^55^ and reconstituted systems^20^ have demonstrated that local laser ablation of the actin cortex can trigger bleb formation. However, the key molecular parameters within the actin cortex that govern bleb initiation remain unclear, because laser ablation disrupts the actin network nonspecifically. In this study, we have identified both *in vitro* and *in silico* that the actin-membrane coupling strength, *R*_C_, and the network connectivity, *R*_X_, are the key parameters regulating bleb formation and the initiation mechanism. Using the *in silico* model, we tested whether external perturbation, either on *R*_C_ or *R*_X_, can induce bleb formation in a specified position. Specifically, we locally reduced *R*_C_ (**Fig. 8a,b**) or *R*_X_ (**Fig. 8c,d**) to different extents under the reference condition where bleb formation via “Rupture” was observed. When the perturbation level was low, a bleb was still formed at random positions. When the perturbation level was sufficiently high, the bleb was consistently formed from the perturbed region. Interestingly, we could switch the mechanism from “Rupture” to “Detachment” by inducing decoupling of a large portion of the cortex from the membrane (**Fig. 8a**). These simulations clarified that the location and mechanism of bleb formation can be controlled by locally manipulating one of the two parameters.

**Figure 8.**
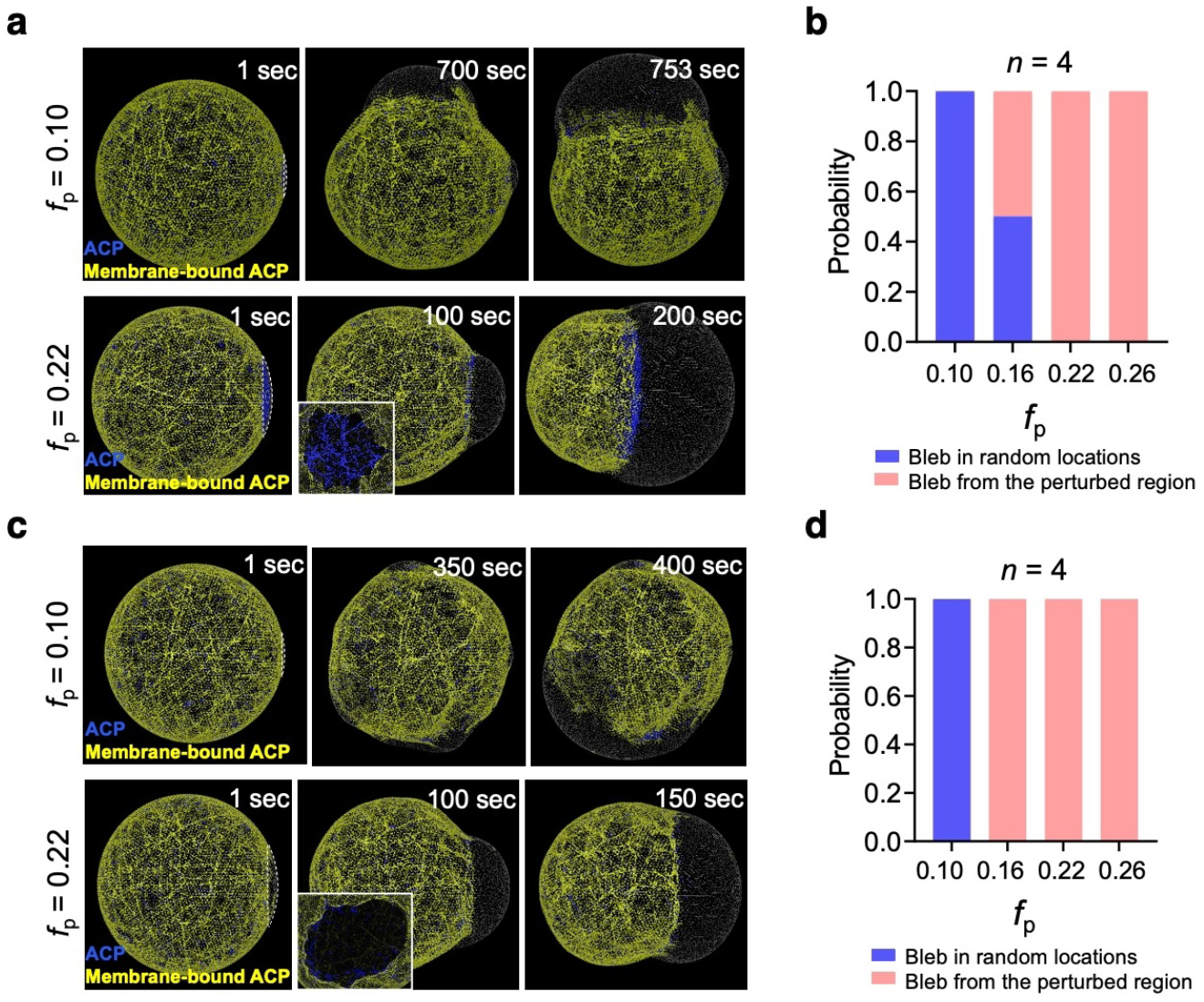
Local perturbation on actin-membrane coupling strength or network connectivity controls bleb position. **a,b**, Perturbation on actin-membrane coupling strength, *R*_C_. **c,d**, Perturbation on network connectivity, *R*_X_. We tested whether a local perturbation could initiate bleb formation from a specified location by selectively preventing ACPs from binding to the membrane (*R*_C_ = 0) or binding to F-actin (*R*_X_ = 0) within a region defined by a fraction of the membrane surface, *f*_p_ (white dashed lines in **a** and **c**; *t* = 1 sec), where *f*_p_ indicates ratio of the perturbed membrane area to the total membrane area. **a**, With *f*_p_ = 0.10 (top), a bleb formed in a different location. With *f*_p_ = 0.22 (bottom), a bleb emerged from the perturbed region, initiated by the detachment of the network from the membrane, as evidenced by the intact network remaining beneath the bleb (inset). **c**, With *f*_p_ = 0.10 (top), a bleb formed in a different location. With *f*_p_ = 0.22 (bottom), a bleb emerged from the perturbed region, initiated by the rupture of the network, as evidenced by a hole in the network beneath the bleb (inset). **b,d**, Dependence of *f*_p_ on the probability of bleb formation from the perturbed region. At *f*_p_ = 0.16, perturbation on *R*_C_ controlled the bleb position with 50% efficiency (**b**), whereas perturbation on *R*_X_ controlled the bleb position with 100% efficiency (**d**). This difference arises because ACPs contribute to both actin-membrane coupling and network connectivity. In the absence of ACPs (*R*_X_ = 0), the network becomes mechanically weaker, making the system more susceptible to bleb formation.

## DISCUSSION

The actin cortex is a ubiquitous structure found in various cells. However, the interaction between actomyosin machinery and the upstream/downstream signaling pathways, which are different between cell types and species, makes it difficult to isolate the contribution of actomyosin machinery to the morphological transition process. In this study, we combined an *in vitro* and *in silico* systems (**Fig. 1, Fig. 6**) to dissect how and to what extent actomyosin machinery contributes to the regulation of membrane deformation without any biochemical signaling cue.

We demonstrated that the single parameter, His-α-actinin concentration *C*_H_, can switch the morphological transition mode of cell-sized liposomes (**Fig. 2, Fig. 5, Fig. 7e-l**). Interestingly, with high actin-membrane coupling strength and high network connectivity with the 3D network organization, the minimal model system in both experiments and simulations always formed a single bleb (**Fig. 5d, Fig. 7l**). This result indicates that the actin cytoskeleton can form cell-scale polarity solely by the mechanics of cytoskeletal proteins and the plasma membrane, even in the absence of spatial biochemical cues. At the initial stage of cell migration mediated by the actin cortex, the cell spontaneously establishes cell-scale polarity to define the front- and rear-end by changing uniform actin cortex to a polarized actomyosin structure^3,5^. Similar polarization is observed in the one-cell stage of *C. elegans* embryos, in which asymmetric actin network contraction segregates PAR polarity proteins to define the anterior-posterior axis^55^. Our findings suggest that the cell tunes the actin network connectivity and cortex-membrane attachment to ensure the robust formation of cell polarity.

There is an ongoing debate about the bleb initiation mechanism^2^. Observations in various cells suggest that the bleb initiation process may involve two distinct mechanisms: the local rupture of the actin cortex^13,57^ or detachment of the membrane from the actin cortex^58^. The mechanism seems to depend on the cell types and animal species, and thus a comprehensive understanding of the two distinct bleb initiation mechanisms is still missing. In particular, it remains unclear which specific parameters determine the switch between these two mechanisms. Here, we demonstrated both *in vitro* and *in silico* that the two mechanisms can be switched by varying the actin-membrane coupling strength *R*_C_ and network connectivity *R*_X_ (**Fig. 4, Fig. 7a-d**). The physical model (**Fig. 4c-e**) in combination with simulations (**Fig. 6, Fig. 7a-d**) clarified that the bleb initiation mechanism depends on whether adhesion between the actin cortex and the membrane is sufficient to sustain myosin-induced forces until F-actin in the network undergoes catastrophic rupture severing by strong tensile forces.

Control of the bleb initiation mechanism is important for regulating cell functions. Recent observations on cancer cells^59^ and primordial germ cells^60^ have revealed that the endoplasmic reticulum (ER) network in the cytoplasm flows into the expanding bleb. This ER network entry stimulates store-operated calcium entry (SOCE)-mediated calcium flux to the expanding bleb through the formation of the STIM/Orai complex between the ER and the plasma membrane. When the bleb is initiated by the detachment mechanism, the remaining cortex may serve as a physical barrier that blocks ER network entry into the expanding bleb. During bleb-based motility, the remaining cortex can prevent the nucleus and other organelles from being translocated to the leading edge. In this study, we showed that the rupture mechanism is preferred in the presence of strong actin-membrane coupling. Previously, we demonstrated using cell-extract-encapsulated droplets that strong actin-membrane coupling is the key factor for transmitting the force generated by the actomyosin machinery to extracellular environments for motility^61^. Therefore, from a physical viewpoint, it is reasonable for motile cells to select the rupture-initiated bleb formation for efficient force transmission. Thus, the two distinct initiation mechanisms likely have specific biological roles, and tight regulation of the bleb initiation process may be crucial for controlling cell functions.

Control of the bleb position is also important. For example, E-cadherin confines the bleb-forming region to a restricted area at the leading edge of migrating cells by exerting frictional forces to impede the backward flow of actomyosin structures^62^. This localized bleb formation is crucial for maintaining the front-rear polarity and directional persistence during migration. In this study, using simulations, we showed local reduction of actin-membrane coupling strength or network connectivity can induce bleb formation at the perturbation site (**Fig. 8**). These observations imply that cells control bleb positions by locally manipulating one of the two physical parameters by ph;osphorylations^63,64.^

Furthermore, we successfully reconstituted the formation of multiple blebs from a minimal set of cytoskeletal elements. A combination of experiments and simulations revealed the active role of the bulk actin network in the formation of multiple blebs (**Fig. 2, Fig. 5, Fig. 7i-r, Fig. S5**). However, the parameter range for the multiple bleb formation was narrow. Indeed, in *in vitro* experiments, multiple blebbing was not a predominant event even under optimal conditions, compared to the single bleb formation, which was robust and occurred with high probability (**Fig. 2, Fig. 5, Fig. S5**). This suggests that additional factor(s) may be required to enable the robust formation of multiple blebs. In cells, the actin network in the bulk cytoplasm is physically linked with the microtubule^65^ and intermediate filament^66^ networks. Future studies will be needed to examine how these composite network organizations contribute to the regulation of bleb number. In addition, the cytoplasm in living cells is considerably more viscoelastic and heterogeneous than in our reconstituted system^67,68^. We confirmed that the encapsulated buffers were one order of magnitude smaller than the viscosity of cytoplasm^69,70^ (**Materials and Methods**). Therefore, pressure release after bleb initiation is likely to propagate rapidly throughout the liposome^24^, thereby suppressing the formation of additional blebs in our reconstituted model. A key future challenge will be to investigate how the viscoelastic properties and spatial heterogeneity of bulk cytoskeletal networks affect the number and position of blebs both *in vitro* and *in vivo*.

In summary, a combination of *in vitro* reconstitution experiments and agent-based modeling has elucidated the extent to which the mechanical properties of the actomyosin network can control the morphological transitions of cell-sized liposomes, providing physical insights into how the cell tunes actin-membrane coupling strength, network connectivity, and actin distribution to control cell shape and biological functions.

## MATERIALS AND METHODS

### Buffer

All the experiments were performed using A50 buffer (50 mM HEPES-KOH pH 7.6, 50 mM KCl, 5 mM MgCl_2_, 1 mM EGTA)^71,72^,73, unless stated separately.

### Protein preparation

Actin was purified from rabbit white skeletal muscle as previously described^74^, snap-frozen in liquid nitrogen, and stored at −80°C in G buffer (2 mM Tris-HCl, pH 8.0, 0.05 mM CaCl_2_, 2 mM NaN_3_, 0.1 mM ATP, 0.5 mM 2-mercaptoethanol). Unphosphorylated smooth muscle myosin (SMM) was purified from chicken gizzards following our previous protocol^73^ and labeled with Alexa Fluor 488 NHS ester (A20000, Thermo) as needed. The samples were snap-frozen in liquid nitrogen and stored at −80°C in A50 buffer containing 1 mM DTT and 1 mM ATP. Recombinant human ZIP kinase (ZIPK, GST-tagged) was prepared as previously described^73^, snap-frozen in liquid nitrogen, and stored at −80°C in A50 buffer containing 1 mM DTT. Recombinant human α-actinin I (6×His-tagged) was prepared according to our previous report^72^. The His-tag was removed using PreScission protease (27084301, Cytiva) and labeled with Alexa Fluor 488-maleimide (A110254, Thermo) as needed. The samples were snap-frozen in liquid nitrogen and stored at −80°C in A150 buffer (50 mM HEPES-KOH, pH 7.6, 150 mM KCl, 5 mM MgCl_2_, 1 mM EGTA) supplemented with 1 mM 2-mercaptoethanol.

The purity of proteins was confirmed by SDS-PAGE (**Fig. S1a**). The concentrations were determined from UV absorbance, using absorption coefficients: 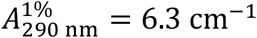 for G-actin, 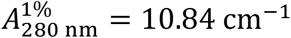 for his-tagged α-actinin I, and 9.61 cm^−^ for his-tag removed α-actinin I, and molecular weights of 42,000 Da for G-actin, 105,300 Da for his-tagged α-actinin I, and 100,000 Da for his-tag removed α-actinin I. The other protein concentrations were determined using Protein Assay Kit (500-0006, Bio-Rad), and molecular weights of 474,000 Da for SMM (hexamer: 2 myosin heavy chains (MHC) + 2 myosin regulatory light chains (MRLC) + 2 myosin essential light chains (MELC)), and 79,084 Da for ZIPK.

### Preparation of lipid-oil mixture

The lipid-oil mixture was prepared as previously described^71^. Briefly, L-α-phosphatidylcholine from chicken egg yolk (egg PC; 840051P, Avanti Polar Lipids), 1,2-dioleoyl-sn-glycero-3-phosphatidylglycerol (DOPG; 840475P, Avanti Polar Lipids), and 1,2-dioleoyl-sn-glycero-3-[(N-(5-amino-1-carboxypentyl)iminodiacetic acid)succinyl] (nickel salt) (DGS-NTA(Ni); 790404C, Avanti Polar Lipids) were mixed at a molar ratio of 85:10:5 and then dissolved in mineral oil (M8410, Sigma-Aldrich) using a bath sonicator at 60°C and the power of 60 W for 90 min. The total lipid concentration in oil was 5 mM.

### Protein encapsulation into liposomes

The inverted emulsion method^75^ was used to encapsulate purified cytoskeletal proteins into cell-sized liposomes^71^. Prior to the experiments, 100-150 μl of the lipid-oil mixture was gently placed on 1 ml of the outer solution (A50 buffer containing 110 mM glucose, 2 mM Trolox, 10 mM DTT, and 0.2 mg ml^-1^ BSA) in a 1.5 ml sample tube, and incubated on ice for >5 min to allow for the assembly of lipid monolayer at the oil/outer-solution interface. Next, 1.5 μl of ×20 E-mix (20 mM ATP, 200 mM phosphocreatine, 2 mg ml^-1^ creatine phosphokinase, 200 mM DTT) in A50 buffer was mixed with 1.5 μl of 10 μM Alexa Fluor 546-conjugated phalloidin (A22283, Thermo) in ×3 A0 + KLC50 buffer (150 mM HEPES-KOH pH 7.6, 50 mM KCl, 15 mM MgCl_2_, 3 mM EGTA), 3 μl of 1 M sucrose and 20 mM Trolox in A50 buffer, 6 μl of 1% (w/v) methylcellulose (1,500 cP; 135-02142, Wako) in A50 buffer, 3 μl of α-actinin I in A150 buffer, and 3 μl of SMM in A50 buffer. Then, 3 μl of ZIPK in A50 buffer was added to initiate myosin phosphorylation. Within a few seconds, this mixture (27 μl) was added to 3 μl of 100 μM G-actin in G buffer and pipetted 5 times to mix instantly. The actin polymerization was initiated by this mixing, and this timing was defined as *t* = 0 sec. Subsequently, 2 μl of the mixture was put into 100 μl of the lipid-oil mixture pre-incubated on ice in a 1.5 ml sample tube, and this sample was vortexed for 10-15 sec to form cell-sized droplets. The emulsion was gently placed on the lipid-oil mixture placed on the outer-solution, and then the tube was centrifuged at 12,000 ×*g* for 60 sec at 4°C to transform droplets into liposomes. In each experiment, the rest of the protein mixture (28 μl) was subjected to the measurement of the osmotic pressure by using a micro-osmometer (Fiske 210, Advanced Instruments). It was confirmed that the pressure difference between the encapsulated protein solution and the outer medium was smaller than ±17 mOsm (less than 5% difference between the inner and outer solutions; the mean difference was -0.23 mOsm, *n* = 222) to reasonably approximate that the liposomes were settled at isotonic condition.

An observation chamber was assembled by placing two double-sided tapes (0.3 mm in thickness) onto a coverslip (60 × 24 mm^2^, thickness No.1, Matsunami) with a coverslip (18 × 18 mm^2^, Matsunami) on top. Immediately after the liposome formation, the liposome solution was perfused into the chamber and sealed with Valap to prevent flow and then warmed to 25 ± 1°C to promote actin polymerization and myosin phosphorylation. The inner surface of the chamber was immediately coated with BSA, which was included in the outer solution, to prohibit non-specific interaction between the liposomes and the coverslips. Note that we added 100 mM of sucrose inside the liposomes and used the equimolar concentration of glucose for the outer solution to create a mass density difference to settle down the liposomes on a bottom coverslip for time-lapse imaging. We confirmed that this mass density difference was enough to sediment the liposomes within 10 min after making an observation chamber and was small enough to keep the liposomes spherical.

The liposomes contained 1 mM ATP, 10 mM phosphocreatine, 0.1 mg ml^-1^ of creatine phosphokinase, 100 mM sucrose, 2 mM Trolox, 10 mM DTT, 0.2% or 0% (w/v) methylcellulose (1,500 cP), 10 μM actin, 0.5 μM Alexa Fluor 546-conjugated phalloidin, 1 μM SMM, and various concentrations of α-actinin I (± his-tag), and 1.4 × 10^−4^ unit ml^-1^ ZIPK. One unit is defined as the amount of ZIPK that phosphorylates 1 μ mol of MRLC in SMM per minute at 25°C. We have reported that 5% (mol/mol) of Alexa Fluor 546-conjugated phalloidin does not change the actin polymerization kinetics, monitored by pyrene-assay^72^.

### Microscopy

Epifluorescence and bright-field images of the liposomes were observed by a custom-built inverted microscope equipped with a ×60 objective (PlanApo NA1.45 oil TIRFM or PlanApo NA1.40 oil, Olympus), an electron-multiplying CCD (charge-coupled device) camera (iXon3 DU-897E-CS0-#BV, Andor Technology). Confocal fluorescence images of the liposomes were observed by an inverted microscope (IX73, Olympus) equipped with a ×60 objective (UPlanSApo NA 1.30 sil, Olympus), a confocal scanner unit (CSU10, Yokogawa), an electron-multiplying CCD camera (iXon3 DU-897E-CS0-#BV, Andor Technology). For both microscopies, a custom-built heat block connected to a water bath (AB-1600, ATTO) was mounted on the objective lens to maintain the sample temperature at 25 ± 1°C.

### Analysis of the distribution of F-actin in liposomes

To quantify the steady-state distribution of F-actin, myosin and ZIPK were excluded from the inner solution of liposomes to eliminate the membrane deformation and contraction of actomyosin networks. The liposomes were observed after >20 min from protein encapsulation, in which time scale it was reasonable to assume a steady state with a balance between polymerization and depolymerization reactions, considering net actin polymerization was almost terminated (**Fig. 1c**). Using confocal microscopy, we measured F-actin density beneath the membrane and compared the value with F-actin density in the bulk region of the liposome (**Fig. 2b,c, Fig. 3b,c, Fig. 5b,c, Fig. S2b-e**).

### Observation of myosin filaments and the length analysis

One micromolar of unphosphorylated SMM labeled with Alexa Fluor 488 (10% labeling ratio) was incubated with 1.4 × 10^−4^ unit ml^-1^ ZIPK in A50 buffer containing 100 mM sucrose, 2 mM Trolox, 0.05 mg ml^-1^ BSA, 1 mM ATP, 10 mM phosphocreatine, 0.1 mg ml^-1^ creatine phosphokinase, and 10 mM DTT at 25°C (ThermoMixer C, Eppendorf) for 120 min in a 1.5 ml sample tube to induce mini-filament formation. BSA was included in the reaction mixture to prevent non-specific adsorption of ZIPK to the tube surface. After the incubation, the sample was immediately diluted 100-fold with the same buffer without ZIPK and BSA. A 4 μl aliquot of the diluted sample was placed onto a coverslip (60 × 24 mm^2^, thickness No.1, Matsunami), covered with a smaller coverslip (18 × 18 mm^2^, Matsunami), and sealed with Valap. The sample was observed using a custom-built total internal reflection fluorescence (TIRF) microscope equipped with a ×60 objective (UPlanApo NA 1.50 oil, Olympus) and EM-CCD camera (iXon3 DU-897E-CS0-#BV, Andor Technology) (**Fig. 1e**). Filament length was measured using fluorescence images (**Fig. S1b**). After background subtraction using a rolling-ball algorithm, binary images were generated, and the length of the major axis of each particle was measured. Overlapping filaments were manually excluded from the analysis.

### Analysis of the phosphorylation rate of myosin

The phosphorylation rate of SMM by ZIPK was measured using urea/glycerol PAGE^35^. One micromolar of unphosphorylated SMM was incubated with 1.4 × 10^−4^ unit ml^-1^ or 0.53 × 10^−4^ unit ml^-1^ ZIPK in A50 buffer containing 100 mM sucrose, 2 mM Trolox, 0.05 mg ml^-1^ BSA, 1 mM ATP, 10 mM phosphocreatine, 0.1 mg ml^-1^ creatine phosphokinase, and 10 mM DTT at 25°C (ThermoMixer C, Eppendorf) for 120 min (1.4 × 10^−4^ unit ml^-1^ ZIPK) or 240 min (0.53 × 10^−4^ unit ml^-1^ ZIPK) in a 1.5 ml sample tube to induce phosphorylation. BSA was included in the reaction mixture to prevent non-specific adsorption of ZIPK to the tube surface. The initial sample volume was 400 μl. Every 10, 20, or 40 min incubation, 30 μl of the reaction mixture was taken from the sample tube and the reaction was terminated by the addition of 1.1 g ml^-1^ urea. Unphosphorylated (0P), monophosphorylated (1P), and diphosphorylated (2P) myosin regulatory light chains (MRLCs) were separated by urea/glycerol PAGE (10 mA, 230 min). The gel was stained using the SYPRO Ruby protein gel stain (S12000, Invitrogen), and the band intensities were analyzed using Image Lab (Bio-Rad) (**Fig. 1c,d, Fig. S4b,c**).

### Analysis of the polymerization rate of actin

The polymerization rate of actin was measured using a pyrene assay^34,72^. Briefly, 8 μl of 100 μM G-actin (5% pyrene-labeled) in G buffer was put into a 384-well microplate (242764, Thremo), then 72 μl of A50 buffer containing ATP, phosphocreatine, creatine phosphokinase, sucrose, Trolox, and DTT was added and pipetted 5 times. Then, the fluorescence intensity was measured using a microplate reader (Fluoroskan FL, Thermo) (**Fig. 1c**). The final concentrations were 10 μM actin, 1 mM ATP, 10 mM phosphocreatine, 0.1 mg ml^-1^ of creatine phosphokinase, 100 mM sucrose, 2 mM Trolox, and 10 mM DTT.

### Analysis of the binding affinity of His-α-actinin to F-actin

To measure the binding affinity of His-α-actinin to F-actin, a co-sedimentation assay was performed^76^. Various concentrations of His-α-actinin in A50 buffer were mixed with 10 μM of G-actin on ice, incubated at 25°C for 60 min in a water bath (AB-1600, ATTO) to polymerize actin, then centrifuged at 100,000 ×*g* for 10 min at 25°C. Identical aliquots of the supernatant (S) and the pellet suspended in A50 buffer with the initial volume before centrifugation (P) were applied to SDS-PAGE. (5-20% gradient precast gel, ATTO), and the gel was stained by Coomassie Brilliant Blue (**Fig. S1c**). The fractions of bound His-α-actinin and free His-α-actinin were calculated from the band intensities^76^. The dissociation constant was determined from the model fitting^77^ (**Fig. S1d**).

### Measurement of the buffer viscosity

The buffer viscosities were measured with Ostwald’s viscometer (*φ* = 0.75 mm; CL2370-02-10, Climbing). The measured viscosities at 24°C were 0.92 mPa s for A50 buffer and 2.27 mPa s for A50 buffer containing 0.2% (w/v) methylcellulose, both of which were one order of magnitude smaller than the viscosity of cytoplasm^69,70.^

### Agent-based model

We developed an agent-based model for a cell-like structure consisting of an actin network and a membrane. The actin network consists of F-actin, actin cross-linking proteins (ACPs), and motors, all of which are simplified via cylindrical segments. The membrane is simplified into a triangulated mesh. The displacements of all points defining the segments and the triangulated mesh are updated at each time step via the Langevin equation with consideration of extensional, bending, and repulsive forces. F-actin is assembled via nucleation and polymerization events but do not undergo depolymerization. However, F-actin can be severed into two filaments if they experience a tensile force beyond a threshold level. ACPs connect pairs of F-actins to form cross-linking points, and motors bind to F-actin and then walk toward its barbed end. At the beginning of each simulation, an actin network is created via self-assembly and interactions of cytoskeletal elements with specified cross-linking density (*R*_X_) and motor density (*R*_M_) within a spherical membrane. In the 2D network simulations, a thin actin network is assembled only beneath a spherical membrane to create a cortical network (**Fig. 6b**, left). In the 3D network simulations, a network is assembled in an entire space within the membrane to create a 3D actin structure (**Fig. 6b**, right). During the network assembly, the spherical membrane does not undergo deformation, and the network and membrane are coupled via a fraction of ACPs defined by the extent of coupling. The fraction is varied between 0 and 1, and it is equal to *R*_C_/*R*_X_, where *R*_C_ is actin-membrane coupling strength. Only after the network assembly, motors start walking to generate mechanical forces, and the membrane also starts deforming by thermal fluctuation and motor-generated forces. The details of the agent-based model and parameters used in simulations are explained in **Supplementary Note 2**.

### Analysis of the orientational angles of motors in simulations

The angle between motor backbones and the circumferential direction was calculated as follows. First, an acute angle between a vector defined by the two ends of the motor backbones, 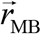, and a vector defined from the membrane center to the centroid of the motor backbones 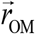, was calculated. Then, the angle was subtracted from 90°:

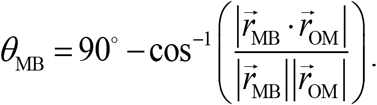

## Data analysis

Image analysis was performed using Fiji (NIH) and MATLAB (MathWorks). Statistical analysis was performed using MATLAB (MathWorks).

## Data Availability

Data are available from the corresponding authors upon reasonable request.

## Author Contributions

M.M. and T.K. designed the research. M.M. prepared proteins, performed *in vitro* experiments, and analyzed the experimental data. M.M. developed the theoretical model. T.K. developed the computational model. F.S.L. performed computer simulations and analyzed the simulation data. All the authors discussed the results and wrote the manuscript.

## Conflicts of Interest

The authors declare that they have no conflicts of interest with the contents of this article.

## Acknowledgments

A plasmid encoding full-length human ZIPK (pGEX-5X-hZIPK) is a kind gift from Kozue Hamao (Hiroshima University, Japan). We thank Yosuke Yamazaki for reviewing the manuscript and providing critical comments, Masatoshi Ichikawa and Kei Yamamoto for discussions, Hibiki Sakata, Mayu Yamaguchi, Takaya Izumi, Tomoko Yamanaka, and Kanako Gomi for their assistance with the experiments, and Nobuko Nishikawa, Terumi Kaji, and Yuki Saito for laboratory management. This work was partly supported by JST PRESTO, Japan (grant no. JPMJPR20ED to M.M.), Grant-in-Aid for Transformative Research Areas (A) from the Ministry of Education, Culture, Sports, Science, and Technology, Japan (grant no. 22H05171 to M.M.), Kishimoto Foundation Research Grant (to M.M.), SPIRITS 2020 of Kyoto University (to M.M.), the Hakubi project of Kyoto University (to M.M.), and the National Institutes of Health (grant no. 1R01GM151628 to F.S.L. and T.K.), and EMBRIO Institute, contract #2120200, a National Science Foundation (NSF) Biology Integration Institute (to T.K.).

## SUPPLEMENTARY FIGURES

**Figure S1.**
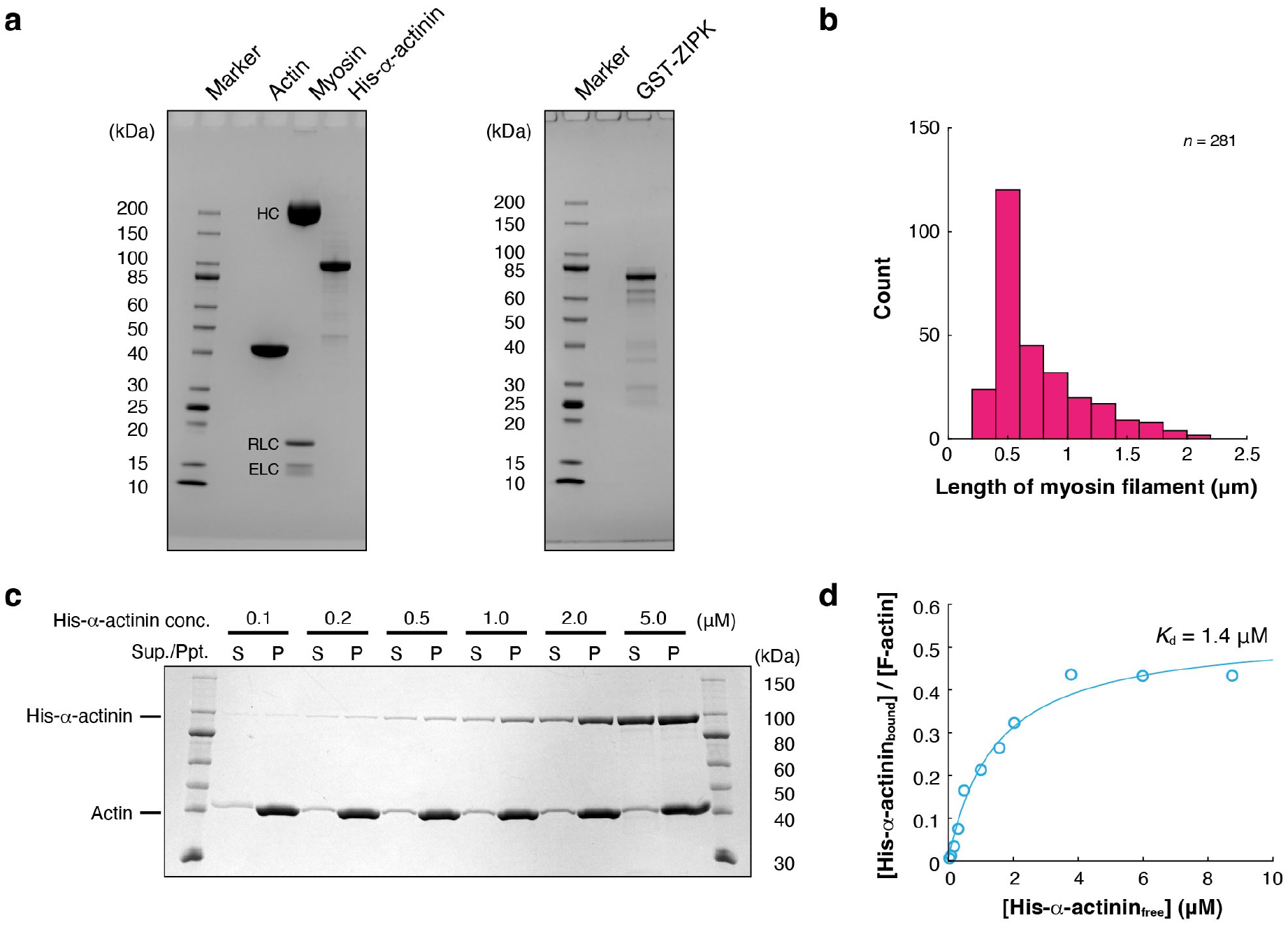
Biochemical properties of proteins used in reconstitution experiments. **a**, Protein purity confirmed by SDS-PAGE. Cytoskeletal proteins (left) and ZIPK (right) used in the experiments were applied to 5-20% gradient polyacrylamide gel. The gels were stained by Coomassie Brilliant Blue. Smooth muscle myosin is a protein complex composed of heavy chain (HC), regulatory light chain (RLC), and essential light chain (ELC). **b**, Length distribution of myosin filaments. The fluorescence images of individual myosin filaments were analyzed. Submicrometer-long mini-filaments (0.64 ± 0.30 μm (mean ± SD, *n* = 550)) were formed by phosphorylation. **c,d**, Dissociation constant *K*_d_ of α-actinin estimated from co-sedimentation assay. **c**, Representative image of SDS-PAGE. Various concentrations of His-α-actinin were mixed with G-actin on ice, incubated at 25°C to polymerize actin, and then centrifuged to precipitate His-α-actinin-F-actin complex. Identical aliquots of the supernatant (S) and the pellet suspended in A50 buffer with the initial volume before centrifugation (P) were applied to 5-20% gradient polyacrylamide gel. The fractions of bound His-α-actinin and free His-α-actinin were calculated from the band intensities. **d**, Experimental data (circles) and the model (solid line): *y* = *B*_max_ *x* / (*K*_d_ + *x*). The dissociation constant *K*_d_ = 1.4 μM was determined from the model fitting using *K*_d_ and *B*_max_ as the fitting parameters (*B*_max_ = 0.53).

**Figure S2.**
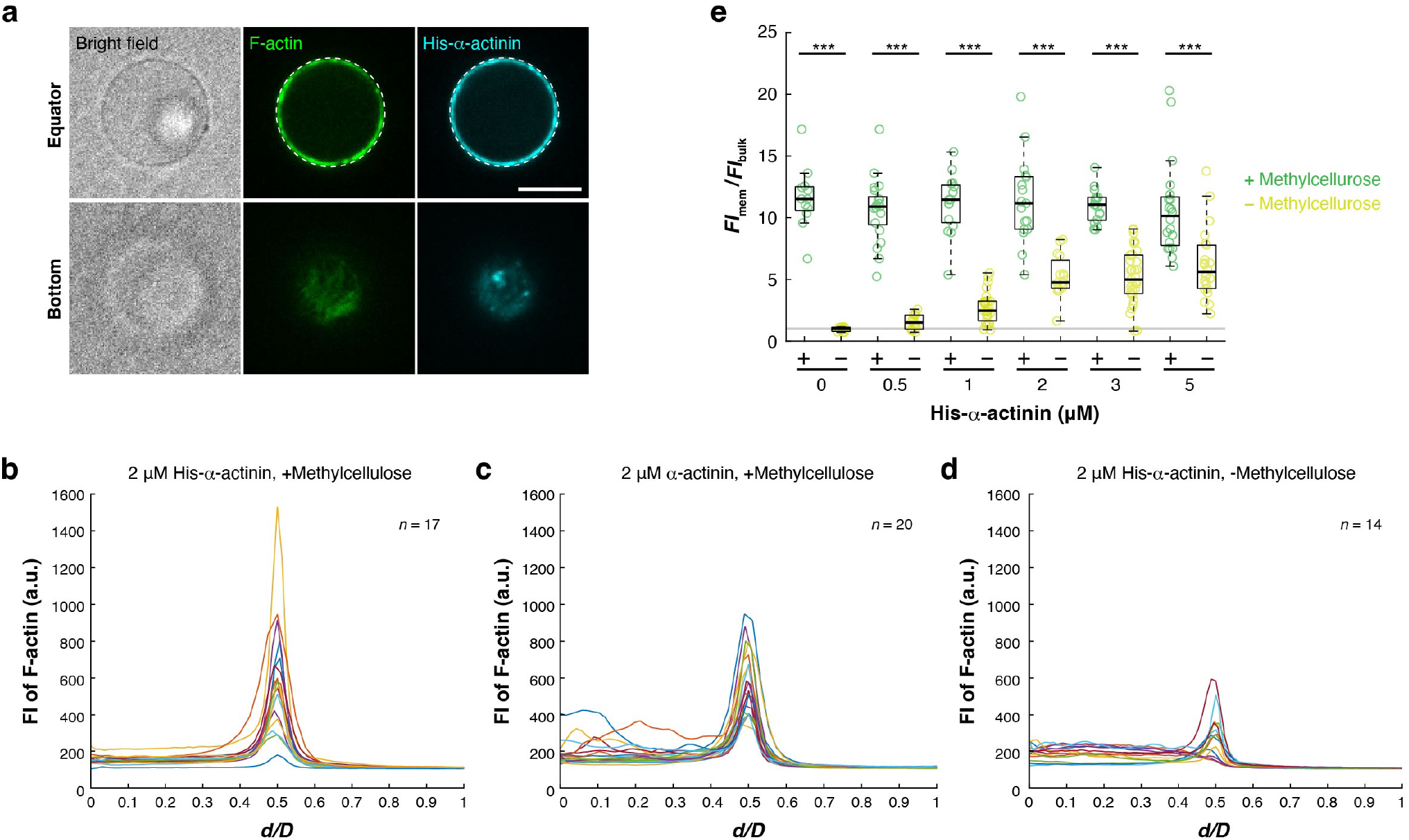
Distribution of proteins in liposomes. **a**, A cross-sectional confocal image of a liposome containing 10 μM actin, 2 μM His-α-actinin (Alexa488-labeled), and methylcellulose at the equator. F-actin and His-α-actinin co-localized beneath the membrane, forming a 2D cortical network. F-actin was visualized using 5% (mol/mol) of Alexa546-phalloidin. The liposome did not contain myosin and ZIPK. White dashed lines indicate the liposome periphery. Scale bar, 10 μm. **b-d**, Fluorescence intensity profiles of F-actin along the radial direction of individual liposomes. Different colors represent different liposomes. **b**, 2 μM His-α-actinin in the presence of methylcellulose. **c**, 2 μM α-actinin in the presence of methylcellulose. **d**, His-α-actinin in the absence of methylcellulose. These are representative raw data from **Fig. 2c, Fig. 3c**, and **Fig. 5c. e**, Comparison of cortical F-actin densities between the presence and absence of methylcellulose. ***: *p* < 0.001 (Welch’s *t*-test, two-sided).

**Figure S3.**
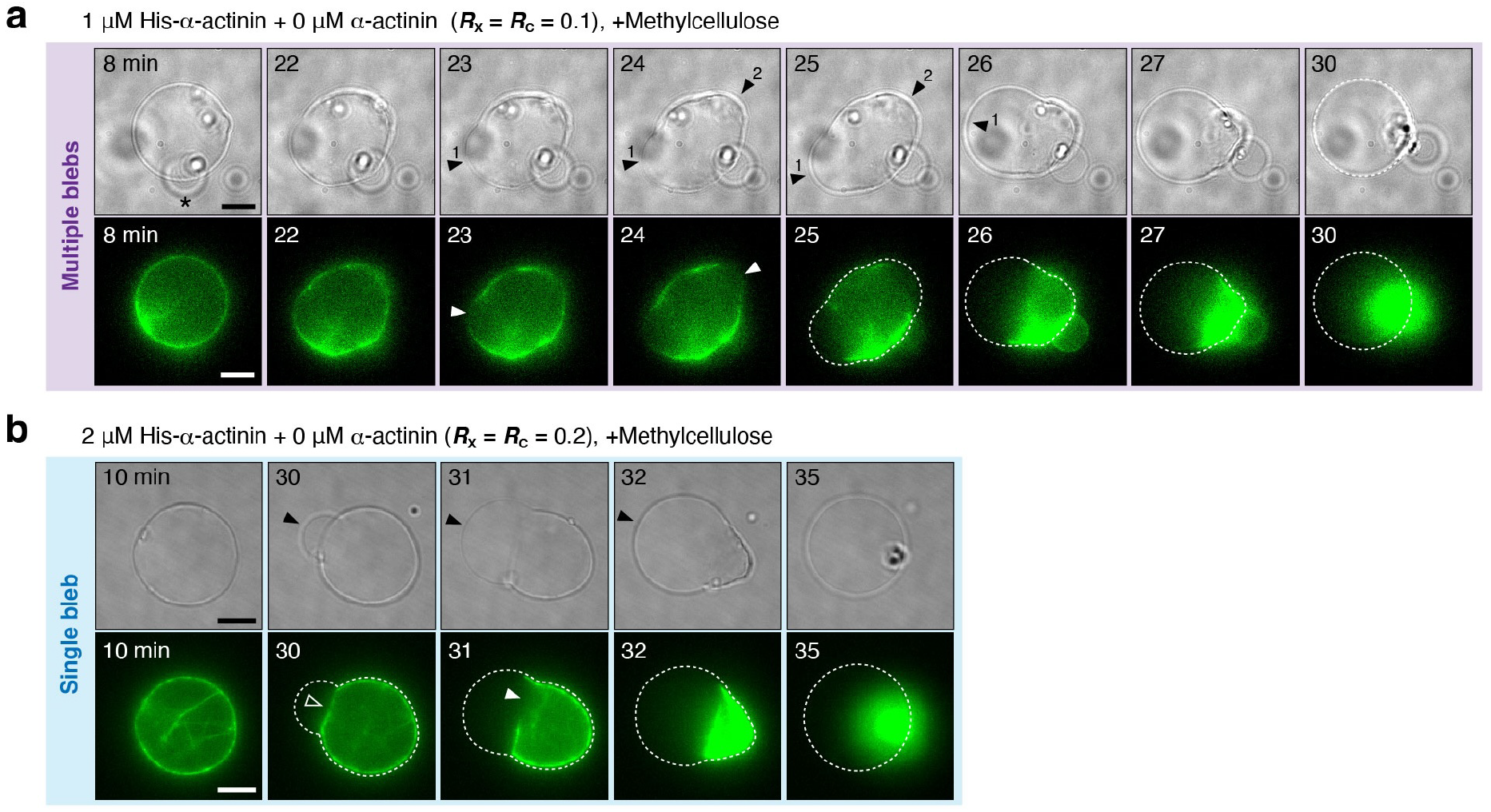
Time-lapse images of liposomes under various conditions. **a**, Time-lapse epifluorescence images of the liposome showing double bleb formation (black arrowheads) in the presence of 1 μM His-α-actinin and methylcellulose (**Movie S5**). The asterisk indicates another small liposome attached to the liposome of interest. White arrowheads indicate the rupture points of the actin cortex. **b**, Time-lapse epifluorescence images of the liposome showing single bleb formation (black arrowheads) by the detachment mechanism in the presence of 2 μM His-α-actinin and methylcellulose. After the detachment (white open arrowhead), the cortex was ruptured (white filled arrowhead) (**Movie S8**). This probability was varied between the concentrations of His-α-actinin (*C*_H_) and α-actinin (*C*_N_): 50% (2/4) at [*C*_H_, *C*_N_] = [2,0], 50% (4/8) at [*C*_H_, *C*_N_] = [1,1], 33% (1/3) at [5,0], 10% (1/10) at [2,3], and 9% (1/17) at [1,2]. For all microscopic images, dashed lines indicate the liposome periphery. Scale bars, 10 μm.

**Figure S4.**
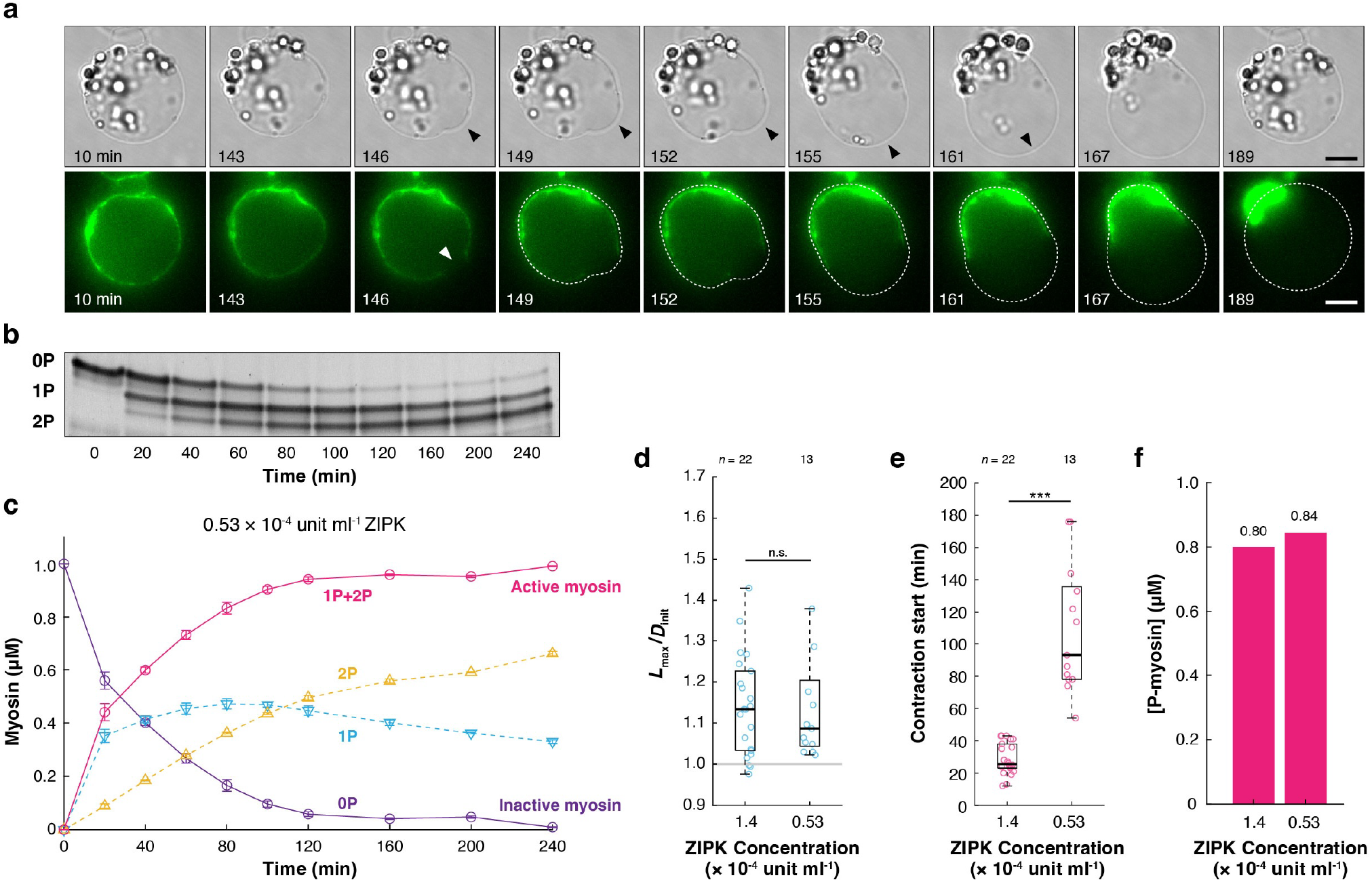
Effects of ZIPK concentration on morphological transition. **a**, Time-lapse epifluorescence images of the liposome containing 10 μM actin, 1 μM SMM, 2 μM His-α-actinin, 0.53 × 10^−4^ unit ml^-1^ ZIPK, and 0.2% (w/v) methylcellulose (1,500 cp) (**Movie S10**). Only ZIPK concentration was different from **Fig. 3e,f**. Black arrowheads and a white arrowhead indicate the blebs and the rupture points of the actin cortex, respectively. White dashed lines indicate the liposome periphery. Scale bars, 10 μm. **b**, The image of urea/glycerol PAGE. 1 μM myosin was incubated with 0.53 × 10^−4^ unit ml^-1^ ZIPK in the presence of 1 mM ATP for the indicated times at 25°C. Unphosphorylated (0P), monophosphorylated (1P), and diphosphorylated (2P) myosins were separated by the gel electrophoresis. **c**, Time courses of the phosphorylation reactions of 1 μM SMM in the presence of 0.53 × 10^−4^ unit ml^-1^ ZIPK measured from the band intensities in **c**. Three independent experiments were performed. Error bars indicate the SDs. **d-f**, Comparison of the morphological transition process of liposomes containing two different concentrations of ZIPK. **d**, Magnitude of membrane deformation. **e**, Contraction start time. **f**, Concentration of phosphorylated myosin predicted from the median values in **e**, and the phosphorylation time courses in **c** and **Fig. 1c**. ***: *p* < 0.001, n.s.: *p* ≥ 0.05 (Welch’s *t*-test, two-sided).

**Figure S5.**
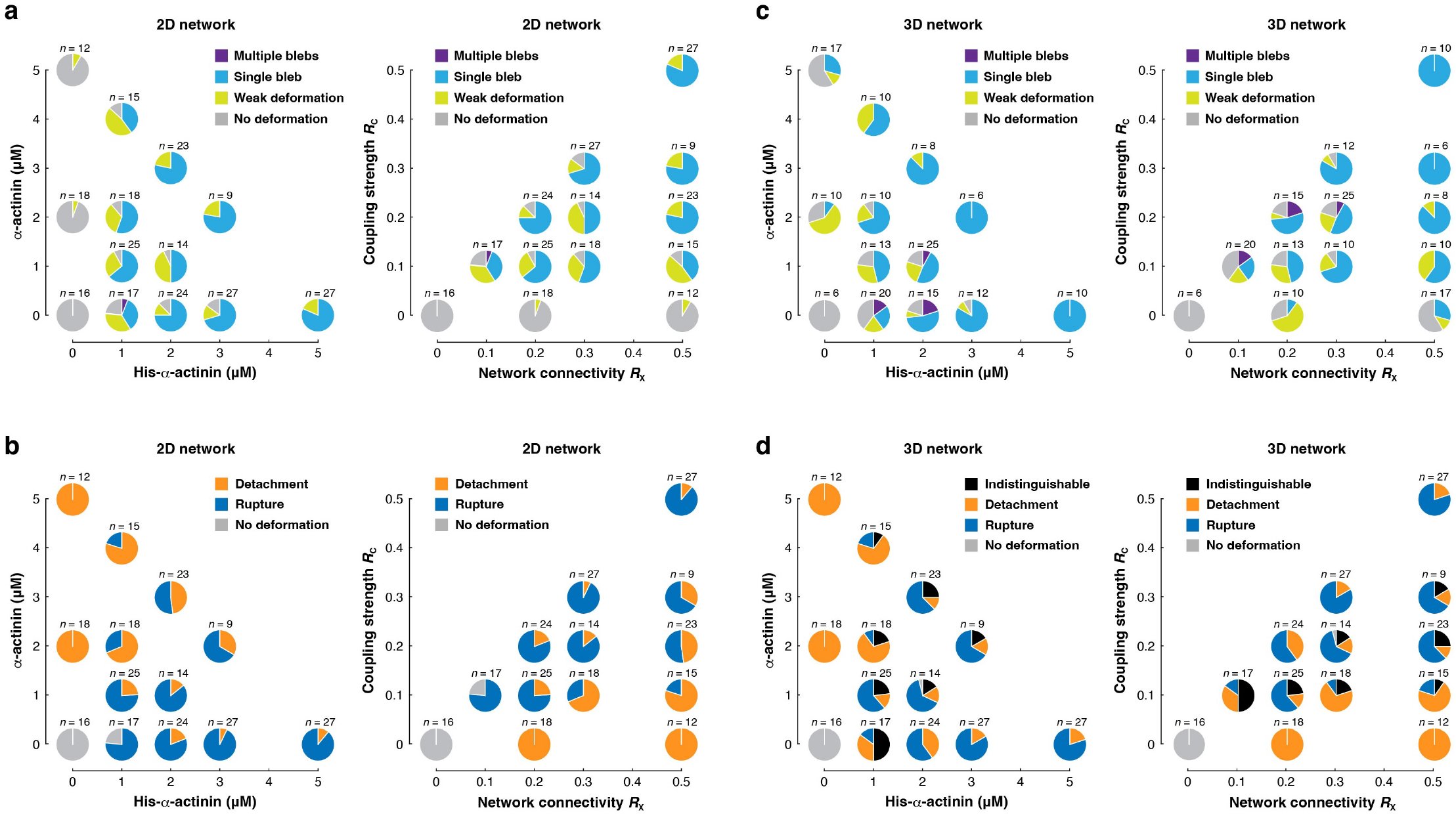
Phase diagram of morphological transition modes and bleb initiation mechanisms in 2D and 3D networks. **a,b**, Phase diagrams of (**a**) morphological transition modes and (**b**) bleb initiation mechanisms in the 2D network conditions (in the presence of methylcellulose), (left) plotted using His-α-actinin concentration (*C*_H_) and α-actinin concentration (*C*_N_) as the parameters, and (right) plotted using the network connectivity (*R*_X_) and actin-membrane coupling strength (*R*_C_) as the parameters. **c,d**, The same types of phase diagrams for the 3D network conditions (in the absence of methylcellulose).

**Figure S6.**
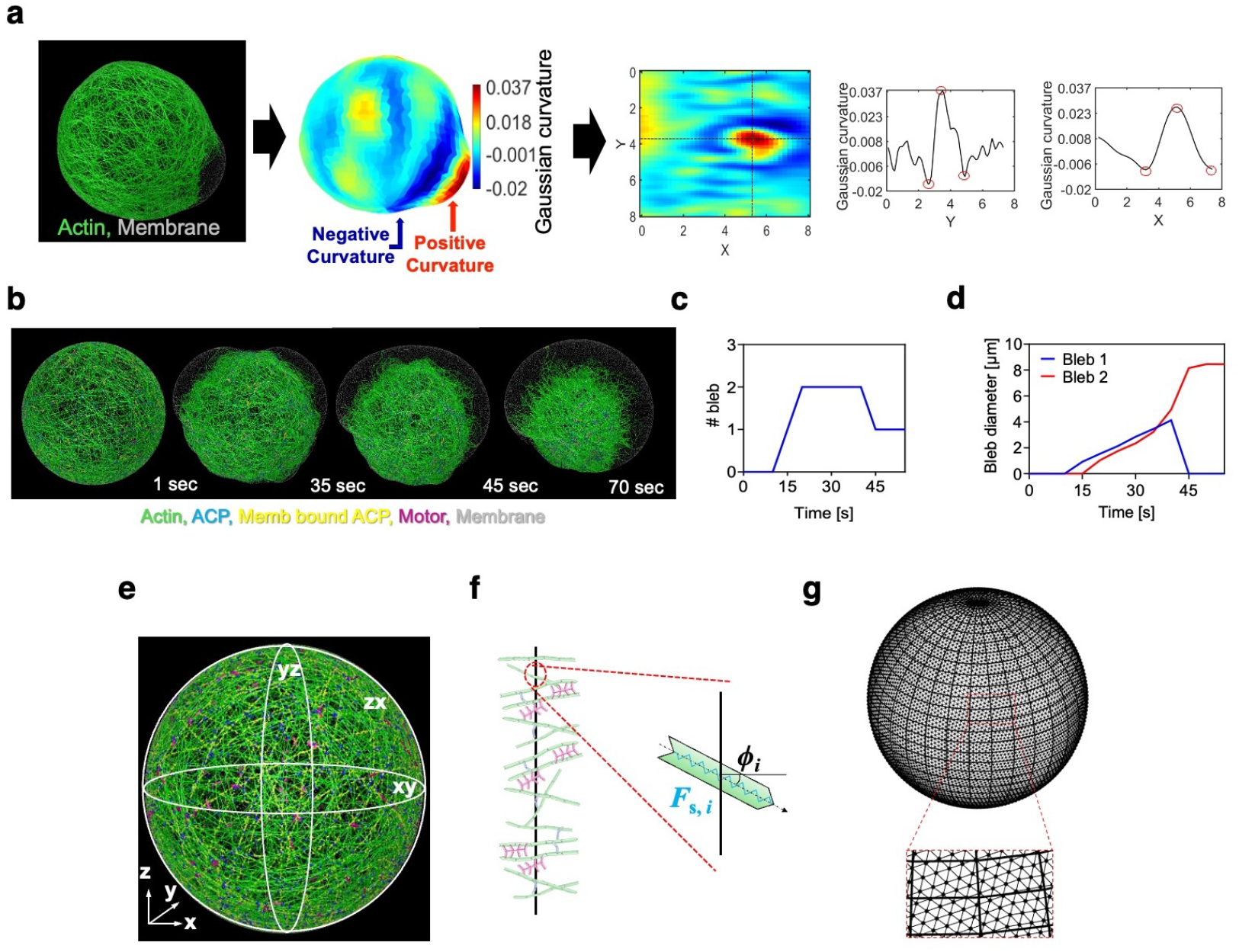
Analysis methods for blebs, global network/membrane tension, and the local density of membrane-bound ACPs. **a**, Detection method for blebs. The curvature of the membrane surface was calculated using the discrete mean curvature approximation based on the Voronoi area and uniform triangular edge length. A region with a positive curvature (red) surrounded by negative-curvature regions (blue) was considered a potential location for a bleb. To estimate bleb radius, the apex of the detected bleb was found, and then the mean distance from the apex to the edges of the bleb where the curvature significantly varies was computed. **b-d**, Representative example of the case showing multiple bleb formation. **b**, Time-lapse images. **c**, Quantification of the number of blebs over time by the detection method shown in **a. d**, The diameter of two blebs changing over time. **e,f**, Analysis method for global network/membrane tension. (**e**) To assess global tension acting on either the actin network or the membrane, we considered *xy, yz*, and *zx* cross-sections that include the instantaneous centroid of the network or the membrane. (**f**) All membrane chains on the triangulated mesh (for membrane tension) or the segments of F-actin, ACPs, and motors (for network tension) crossing each cross-section (indicated by the black line) were identified. The total tension on each cross-section was then determined as the sum of *F*_*s,i*_ cos *ϕ*_*i*_ over all chains or segments. **g**, Analysis method for the local density of membrane-bound ACPs. The membrane surface area was discretized into quadrilateral and triangular surface elements using spherical coordinates. Details of all these analyses are described in **Supplementary Note 2**, Sections 9-11.

**Figure S7.**
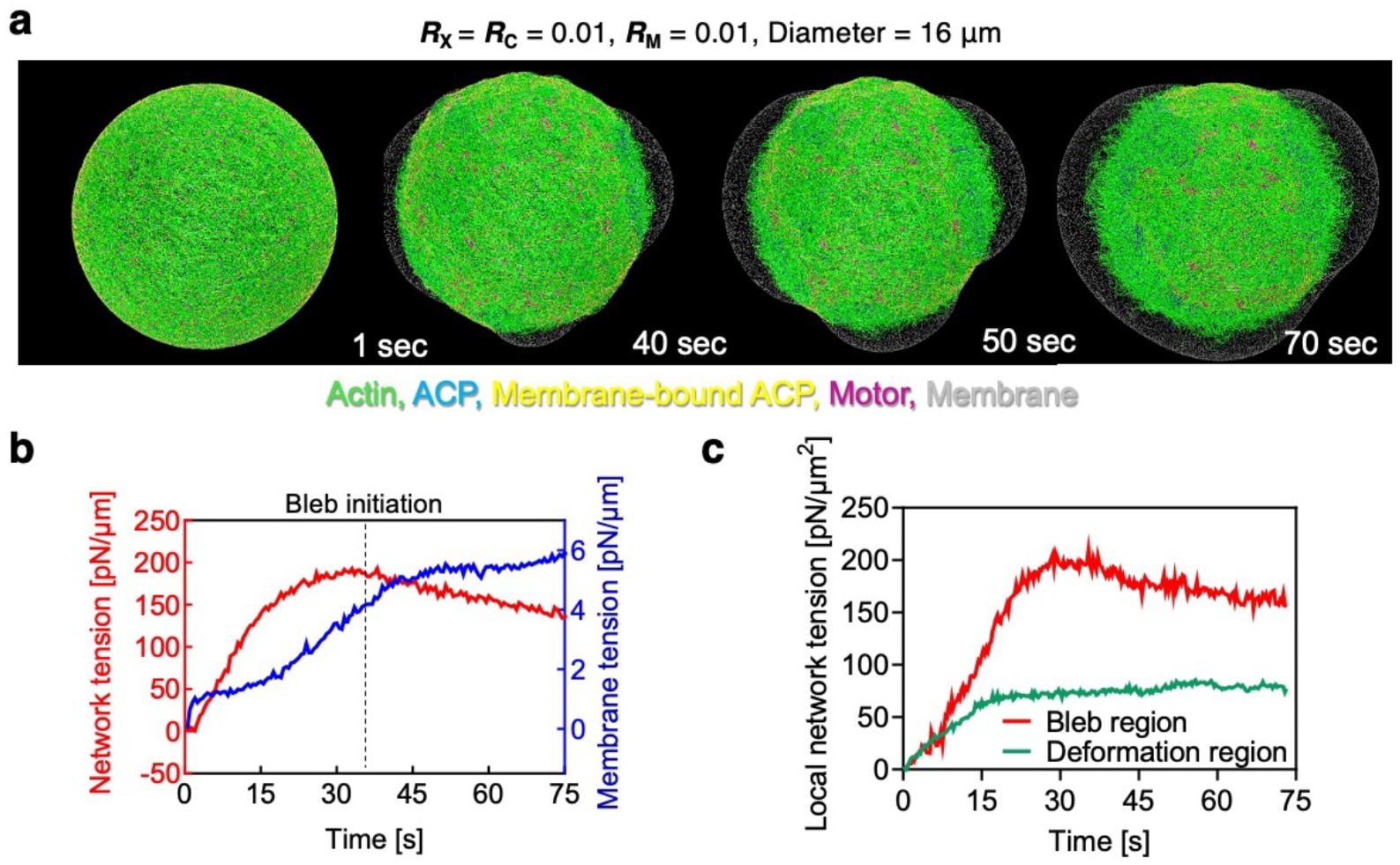
Bleb formation and network/membrane tension observed in a larger vesicle with 16 μm in diameter. **a**, Snapshots of the formation, expansion, and mergence of three blebs (**Movie S15**). **b**, Global tension acting on the network (red) and the membrane (blue) over time. The blebs started emerging from ∼35 s, so the global network tension decreased from ∼35 s. Membrane tension gradually increased from ∼35 s. **c**, Comparison of local network tension between the bleb-forming and deformation-only regions until bleb initiation. Local network tension in the bleb-forming region was noticeably higher than that in the deformation-only region.

**Figure S8.**
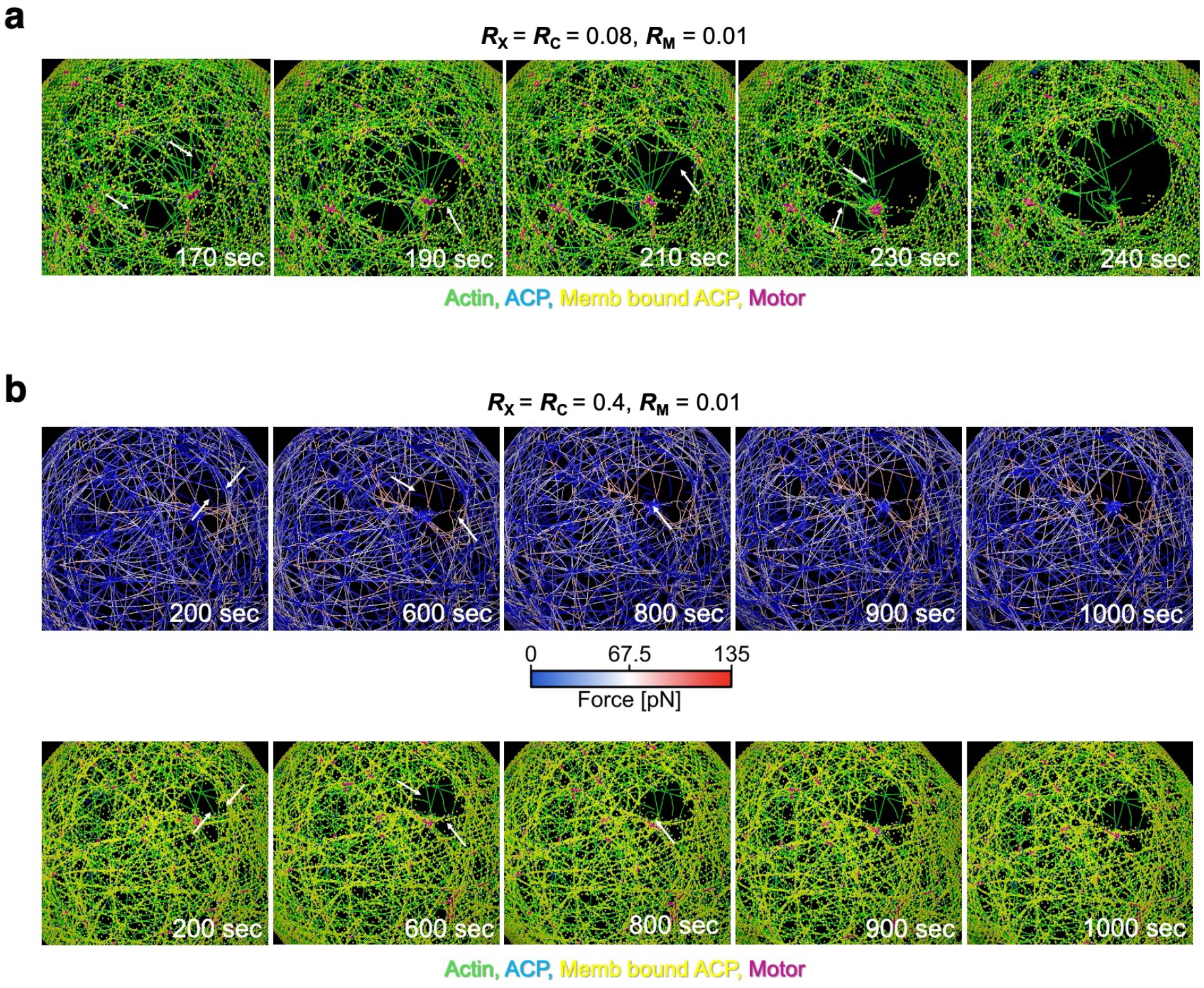
Time evolution of a network rupture induced by F-actin severing. **a**, Intermediate network connectivity and actin-membrane coupling strength (*R*_X_ = *R*_C_ = 0.08). Successive severing events of F-actin indicated by arrows led to the initial formation and expansion of a rupture on a network. **b**, High network connectivity and actin-membrane coupling strength (*R*_X_ = *R*_C_ = 0.4). Several severing events still occurred, but a network rupture was not followed by significant expansion.

**Figure S9.**
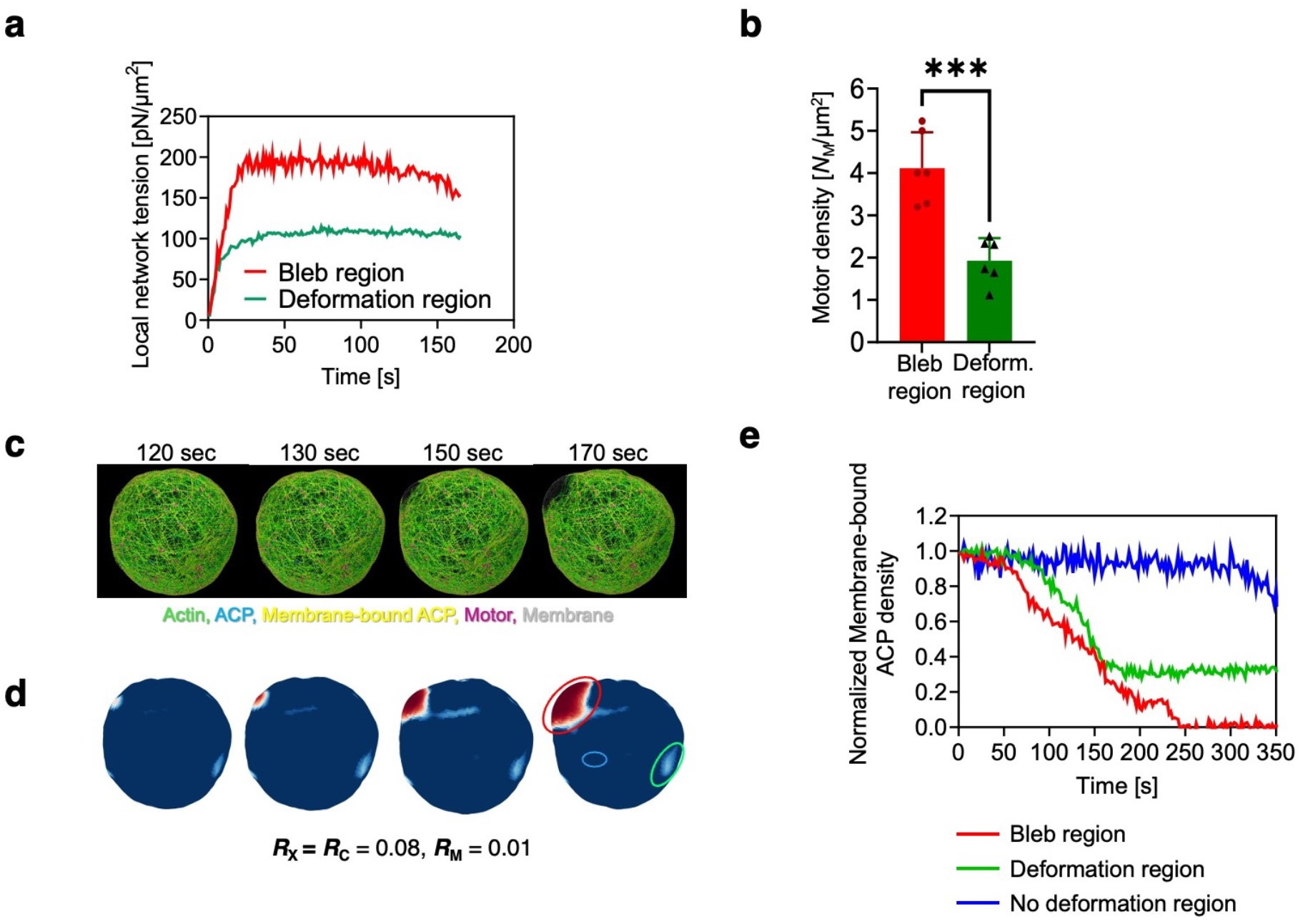
Analysis of local tension, motor density, and actin-membrane coupling. **a**, Comparison of local network tension between the bleb-forming and deformation-only regions developing until bleb formation (The bleb was detected at *t* = ∼150 s in this example). These measurements were performed using the same vesicle. Local network tension measured near the bleb-forming region was substantially higher than that measured in the deformation-only region. **b**, Local motor density measured in the two regions shown in **a**. The density shows a substantial difference, implying that locally higher actomyosin contractility led to a network rupture. ***: *p* < 0.001 (Welch’s *t*-test, two-sided). **c**, Snapshots of the formation of a single bleb at four different time points. **d**, Quantification of the local density of membrane-bound ACPs in the same vesicle at the same time points as **c**. Not all regions with lower density evolved to a bleb. **e**, The local density of membrane-bound ACPs developing over time in bleb-forming (red line), deformation-only (green line), and no-deformation (blue line) regions, as shown in **d** with red, green, and blue circles, respectively. The local density continuously decreased to nearly zero only in the bleb-forming region. The deformation-only region also showed a decrease in the local density, but the local density stopped decreasing and reached a plateau after substantially decreasing, and the network rupture did not expand significantly. The no-deformation region did not show a noticeable decrease in the local density.

**Figure S10.**
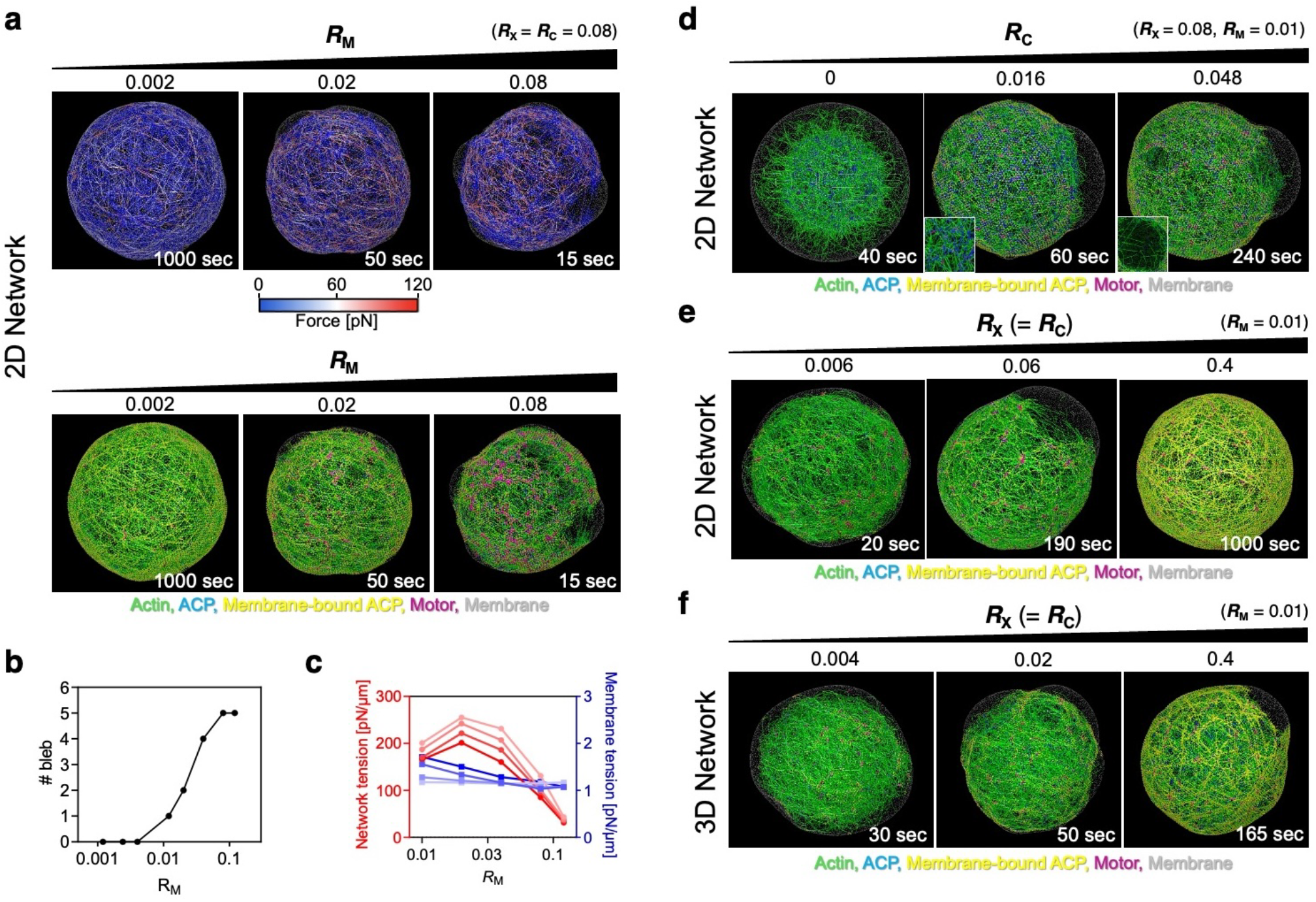
Morphological transitions and bleb formation under different conditions. **a-c**, Effects of the motor density, *R*_M_ (*R*_X_ = *R*_C_ = 0.08). **a**, Representative snapshots of the vesicles with different *R*_M_. (top) Force distribution. (bottom) F-actin, ACPs, motors, and a membrane shown by different colors. ACPs are shown in two different colors, depending on whether or not they are coupled to the membrane. **b**, Number of blebs as a function of *R*_M_. **c**, Network and membrane tension with a variation in *R*_M_. Higher *R*_M_ led to decreased network tension because of impaired connectivity due to several F-actin severing events, whereas membrane tension hardly varied. **d-f**, Snapshots showing the same structures as those shown in **Fig. 7a, e**, and **i**, respectively. Instead of forces, these snapshots show F-actin, ACPs, motors, and a membrane via different colors. ACPs are shown in two different colors, depending on whether or not they are coupled to the membrane.

## SUPPLEMENTARY NOTE 1

In the absence of the hydrostatic pressure (Δ*P =* 0), the actin-bound His-α-actinin density 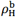 is governed by the kinetic equation:

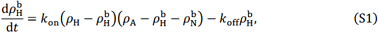

where 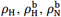, and *ρ*_A_ indicate the surface densities of all His-α-actinin, His-α-actinin bound on F-actin, α-actinin bound on F-actin, and all F-actin beneath the membrane, respectively. *k*_on_ and *k*_off_ are the rate constants for binding and unbinding between His-α-actinin and F-actin, respectively. Considering that Δ*P* tends to separate the membrane from the actin cortex, and assuming that each His-α-actinin-F-actin bond is equally stressed, the force per bond, *f*, becomes 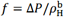. Using Bell’s kinetics model^1^, the kinetic equation (**Eq. S1**) is replaced by

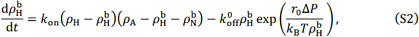

where 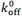 is the zero-force unbinding rate constant, and *r*_0_ is a characteristic distance to overcome the energetic barrier for bond separation^1-3^. If it is assumed that F-actin near the membrane retains many free binding sites for α-actinin 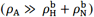, **Eq. S2** can be replaced by

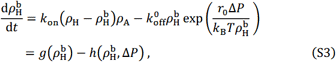

where 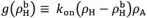 and 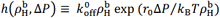 are the binding and unbinding rates, respectively. At the initial condition (*t* = 0), Δ*P* ≃ 0 because forces generated by myosin are negligibly small. If it is assumed that the system is initially at an equilibrium 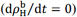, **Eq. S3** becomes

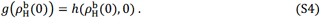

Therefore, we obtain

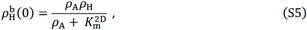

where 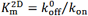 is the dissociation constant of His-α-actinin from F-actin in a 2D network. Subsequently, myosin is gradually phosphorylated and begins to generate contractile forces on the cortical actin network, leading to an increase in Δ*P*. A small increase in Δ*P* slightly elevates the unbinding rate, 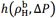, leading to a slight decrease in 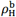, after which the system is expected to reach a new equilibrium 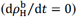. By contrast, when large Δ*P* is applied, the unbinding rate, 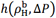, becomes much larger than the binding rate, 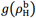, causing 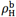 to rapidly approach zero. There will thus exist the critical pressure, Δ*P*^*^, that is sufficient to detach all His-α-actinin from actin filaments.

To evaluate the critical pressure, Δ*P*^*^, we consider the balance between binding and unbinding rates, 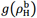 and 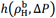, which are plotted in **Fig. SN1** as functions of 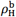. The unbinding rate term (blue lines) contains Δ*P*(*t*) as a parameter. At the initial state (*t* = 0) when myosin-generated force is small enough to be negligible as described above: Δ*P*(0) ≃ 0, both terms are linear and intersect at a single point (**Fig. SN1**, cross), which gives a stable equilibrium 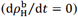. At *t* > 0, Δ*P*(*t*) gradually increases as myosin phosphorylation proceeds, rendering the unbinding rate term nonlinear. Consequently, the two terms have two intersections: a stable fixed point (**Fig. SN1**, filled circle) and an unstable fixed point (**Fig. SN1**, open circle). Since 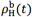 at the stable fixed point is smaller than 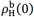 and 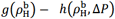 is negative between 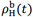 and 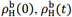 reaches the stable fixed point for each *t* (> 0). A further increase in Δ*P*(*t*) shifts the unbinding rate curve upward, decreasing 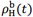 along the red line. When Δ*P*(*t*) reaches Δ*P*^*^, the two terms are tangent, and the stable and unstable fixed points merge at 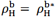. For Δ*P*(*t*) > Δ*P*^*^, no stable fixed point exists, and 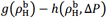 is negative for all 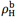. Hence, 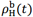 decreases monotonically to zero, eventually leading to the complete detachment of actin filaments from the membrane. Therefore, Δ*P*^*^ is the critical pressure that initiates “Detachment”.

Using the facts that the binding and unbinding rates are equal and their slopes are also equal at the point of tangency, 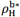 and Δ*P*^*^ satisfy the following equations,

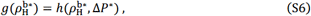

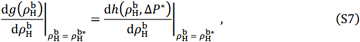

which lead to

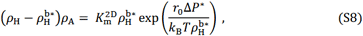

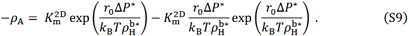

By combining **Eq. S8** and **Eq. S9** and eliminating 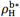, we obtain the following equation:

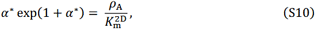

Where

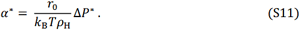

As shown in **Fig. SN2**,

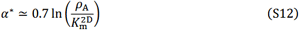

for an extensive range of 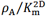. Therefore, by substituting **Eq. S12** to **Eq. S11**, we obtain the critical pressure for “Detachment”:

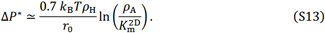

**Figure SN1.**
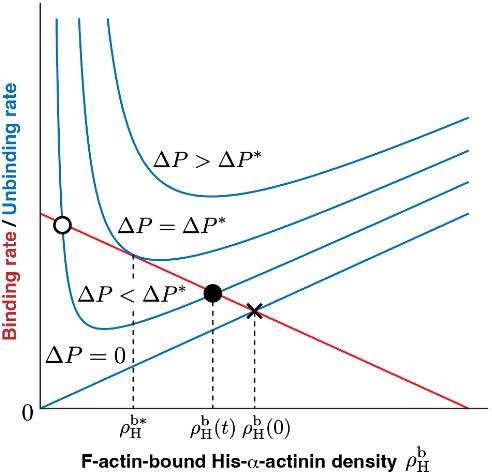
Relationship between the binding and unbinding rates (the first and second terms on the right-hand side of **Eq. S3**, respectively) as a function of F-actin-bound His-α-actinin density, 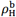. At *t =* 0, the two rates are balanced at a single intersection point (cross). As *t* increases and Δ*P*(*t*) rises, a stable fixed point (filled circle) and an unstable fixed point (open circle) gradually approach each other and merge at the critical pressure, Δ*P*^*^.

**Figure SN2.**
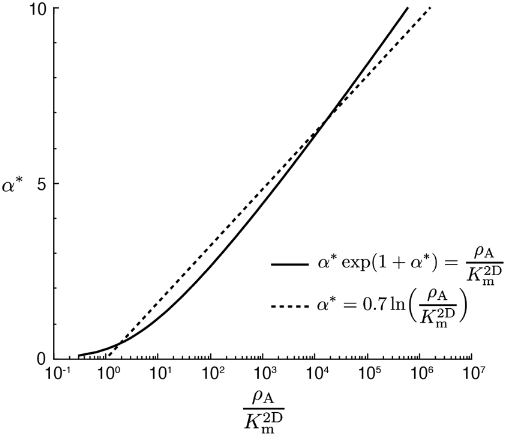
Comparison between **Eq. S10** (solid line) and **Eq. S12** (dashed line).

## SUPPLEMENTARY NOTE 2

### 1. Brownian dynamics simulations via the Langevin equation

In the agent-based model, elements are defined by points (i.e., nodes), and cylindrical segments exist between the points. The displacements of points are governed by the Langevin equation with inertia neglected:

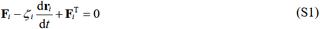

where **r**_*i*_ is the position of the *i*th point, *ζ*_*i*_ is a drag coefficient, *t* is time, **F**_*i*_ is a deterministic force, and 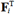 is a stochastic force. The magnitude of 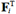 is determined by the fluctuation-dissipation theorem:

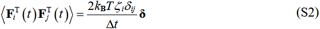

where *δ*_*ij*_ is the Kronecker delta, ***δ*** is a second-order tensor, and Δ*t* is a time step. In most simulations, Δ*t* is 1.15×10^−4^ s.

The drag coefficients of points are computed using an approximated form for a cylindrical object^1^:

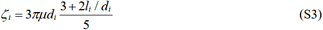

where *μ* is the viscosity of a medium, and *l*_*i*_ and *d*_*i*_ are the length and diameter of a cylindrical segment connected to the point, respectively. The positions of points are updated at each time step using the Euler integration scheme:

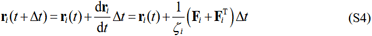

Three types of deterministic forces maintain lengths, distances, and angles involved with components close to their equilibrium values. Extensional forces maintain the equilibrium lengths of cylindrical segments or chains, bending forces maintain angles formed by interconnected segments or dihedral angles, and repulsive forces represent volume-exclusion effects acting between overlapping components. The extensional and bending forces originate from harmonic potentials:

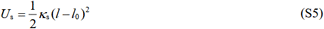

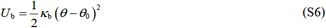

where *κ*_s_ is extensional stiffness, *l* and *l*_0_ are instantaneous and equilibrium lengths, *κ*_b_ is bending stiffness, and *θ* and *θ*_0_ are instantaneous and equilibrium angles. All details of this model are explained below.

### 2. Simplification and mechanics of cytoskeletal components

F-actin is simplified into serially connected cylindrical segments. Each segment has polarity defined by barbed and pointed ends. Extensional stiffness of F-actin (*κ*_s,A_) maintains the length of actin segments close to its equilibrium (*l*_0,A_ = 140 nm). Bending stiffness of F-actin (*κ*_b,A_) maintains an angle formed by two adjacent actin segments close to its equilibrium (*θ*_0,A_ = 0 rad). The persistence length of F-actin (*l*_p_) calculated from *κ*_b,A_ and *k*_B_*T* corresponds to ∼9 μm (ref. 2).

Actin crosslinking proteins (ACPs) consist of two cylindrical segments connected at their center point. Extensional stiffness of ACPs maintains the length of each ACP segment close to its equilibrium (*l*_0,ACP_ = 23.5 nm). Bending stiffness of ACPs (*κ*_b,ACP_) maintains an angle formed by two ACP segments at the center point close its equilibrium (*θ*_0,ACP_ = 0 rad). The geometry of ACPs is analogous to α-actinin^3^.

Each motor has a backbone structure with 7 segments and 8 points. On each point, two motor arms are connected, so the total number of motor arms is 16 (*N*_arm_ = 16). Each arm represents the kinetics and force-velocity relationship of the ensemble consisting of 4 myosin heads (*N*_h_ = 4). Thus, the total number of myosin heads represented by each motor is 64, which is not far from ∼56 myosin heads in a single non-muscle myosin mini-filament^4^. The equilibrium length of each motor backbone segment (*l*_0,M1_ = 42 nm) is regulated by extensional stiffness (*κ*_s,M1_). Thus, the equilibrium length of a whole motor backbone is 294 nm, nearly equal to the length of non-muscle myosin mini-filaments^5^. An equilibrium angle formed by adjacent backbone segments (*θ*_0,M_ = 0 rad) is regulated by bending stiffness (*κ*_b,M_). The extension of each motor arm is regulated by the two-spring model with the stiffnesses of transverse (*κ*_s,M2_) and longitudinal (*κ*_s,M3_) springs. The transverse spring regulates an equilibrium distance (*l*_0,M2_ = 13.5 nm) between the backbone point where the motor arm is connected and an actin segment where the other end of the motor arm is bound, and the longitudinal spring maintains a right angle between the motor arm and the actin segment (*l*_0,M3_ = 0 nm).

Forces exerted on actin segments by bound ACPs and motors are distributed to the barbed and pointed ends of the actin segments as described in our previous work^6^. In the network, repulsive forces are calculated only between actin filaments (among cytoskeletal components), following a harmonic potential^6^:

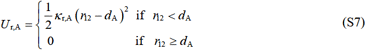

where *κ*_r,A_ is the strength of the repulsive force, *r*_12_ is a minimum distance between two neighboring actin segments, and *d*_A_ is the diameter of the actin segment.

### 3. Simplification and mechanics of the membrane

A cell membrane is simplified into a triangulated mesh with area and volume conservation. Extensional stiffness of the membrane (*κ*_s,MB_) governs the equilibrium length of chains (*l*_0,MB_ = ∼70 nm) constituting the mesh. Bending stiffness of the membrane (*κ*_b,MB_) regulates dihedral angles formed by adjacent triangular elements on the mesh close to their equilibrium value (*θ*_0,MB_ = 0 rad). The area of each triangular element and the entire volume within the membrane are conserved following harmonic potentials:

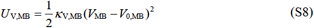

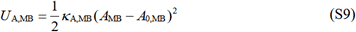

where *κ*_V,MB_ and *κ*_A,MB_ are the strengths of volume and area conservation, *V*_MB_ is volume, *A*_MB_ is area, and the subscript 0 denotes the equilibrium value.

Repulsive forces are applied between triangular elements on the mesh with strength, *κ*_r,MB_, to prevent them from crossing each other. The repulsive forces originate from the following potential:

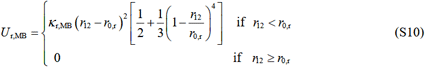

where *κ*_r,MB_ is the strength of the repulsive force, and *r*_0,r_ is a distance below which the repulsive force starts acting. In this case, *r*_0,r_ is equal to the thickness of the membrane mesh (*r*_0,r_ = *d*_MB_). Note that this repulsive force increases more rapidly as the two membrane elements get closer to each other.

### 4. Actin dynamics

The formation of F-actin is initiated from a nucleation event occurring at a constant rate, *k*_n,A_, with the appearance of one cylindrical segment in a random position. The polymerization of F-actin is simulated by adding cylindrical segments at the barbed end of existing filaments at a constant rate, *k*_+,A_. The nucleation and polymerization events are skipped if any part of the new segment is located outside a designated space for network assembly. The depolymerization of F-actin does not take place. The severing of F-actin occurs in a deterministic manner; if a tensile force acting on an actin segment becomes greater than *F*_sev_ = 300 pN (ref. 7), the segment is removed from the filament where it belongs. As a result of each severing event, one filament is divided into two filaments.

### 5. Dynamic behaviors of ACPs

ACP segments bind to binding sites located every 7 nm on actin segments at a constant rate, *k*_+,ACP_ without preference for crosslinking angle. Unlike our previous studies^8^, the unbinding of ACP segments from F-actin is not permitted to allow a network rupture to occur only via F-actin severing. If an actin segment disappears as a result of the F-actin severing, ACPs previously bound to the segment can bind to other F-actin.

Our assumption in the model that the unbinding event of ACPs from F-actin is not allowed may not look reasonable. However, this assumption can be justified as follows. It has been demonstrated *in vitro* that the unbinding force of single skeletal muscle α-actinin from F-actin ranged from 1.4 to 44 pN (18 pN on average) (ref. 9) or 40 to 80 pN (ref. 10), depending on the loading rate. On the other hand, the severing force of F-actin ranges between 200 pN and 600 pN, depending on the magnitude of twisting^7^. Assuming that 40 pN is the mean unbinding force of non-muscle α-actinin I from F-actin and that each α-actinin-F-actin bond is equally stressed, F-actin severing is expected to dominate the network rupture mechanism when there are more than 5-15 α-actinin molecules bound to a single filament. Using the mean length of F-actin in simulations (∼4.2 μm in the 2D network and ∼5.2 μm in the 3D network; **Fig. 6b**, caption) and the size of actin monomers (5.4 nm), the mean number of α-actinin molecules bound to a single actin filament is estimated to be more than 6 molecules under all tested conditions (minimum case: 6 molecules at *R*_X_ = 0.004; reference case: 124 molecules at *R*_X_ = 0.08). In addition, the mean F-actin length in the experiments is estimated to be ∼4.6 μm (ref. 11) (the same actin concentration and buffer were used in this report), which is comparable to the simulations. Therefore, it is reasonable to approximate that severing dominates the network rupture events in the simulations.

### 6. Dynamic behaviors of motors

Motors are created by the self-assembly of backbone segments. Motor arms connected to the backbone points bind to binding sites on actin segments at a constant rate, *k*_+,M_ = 40*N*_h_ s^-1^. The walking (*k*_w,M_) and unbinding (*k*_u,M_) rates of the motor arms are determined rigorously by the parallel cluster model to capture the mechanochemical cycle of non-muscle myosin II^13,14^. The details of implementation and benchmarking of the parallel cluster model in our model were thoroughly described in our previous study^6^. *k*_w,M_ and *k*_u,M_ in the model are lower with a higher applied load, based on an assumption that motors behave as a catch bond. The unloaded walking velocity and stall force of motor arms are set to ∼140 nm/s and ∼5.7 pN, respectively. If an actin segment where motor arms are bound disappears due to the F-actin severing, motor arms can bind to other actin segments.

### 7. Interactions between the cytoskeletal elements and the membrane

Repulsive forces are applied between all cytoskeletal elements and the membrane mesh to keep the cytoskeletal elements within the membrane. The repulsive forces originate from the potential shown in **Eq. S10** and the same strength, *κ*_r,MB_. Here, *r*_0,r_ is equal to the average of membrane thickness and the diameter of a neighboring cytoskeletal element, *r*_0,r_ = (*d*_MB_ + *d*_*i*_)/2, where *i* is either A (actin), ACP, or M (motor).

A fraction of ACPs are selected at the beginning of simulations based on the strength of network-membrane coupling. The selected ACPs can feel an attractive force from the membrane if a distance between the ACPs and the membrane is smaller than *d*_ACP_ + *d*_MB_. The attractive force is defined by the following harmonic potential:

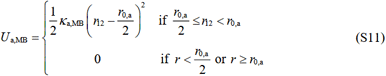

where *κ*_a,MB_ is the strength of the attractive force, and *r*_0,a_ is equal to *d*_ACP_ + *d*_MB_. Note that the attractive force is always applied in a direction normal to the closest triangular mesh element, meaning that ACPs can freely slide along the membrane surface with the maintenance of the equilibrium distance from the membrane. Drag forces acting between ACPs and the membrane are ignored for simplicity, meaning that ACPs bound to the membrane still experience a drag force defined by **Eq. S3**.

In the simulations, it is assumed that the membrane-bound ACPs dissociate from the membrane if a distance between them (*r*_12_) becomes greater than *r*_0,a_. At this threshold distance, ACPs experience ∼30 pN (**Eq. S11** and **Table S1**). In the experiments, α-actinin binds to the membrane through the interaction between the His-tag and Ni-NTA on the lipid. It has been measured by AFM that a dissociation force for His-tag-Ni-NTA interaction is >100 pN (ref. 12), which is larger than the unbinding force of α-actinin from F-actin (1.4-80 pN) (refs. 9,10). Therefore, it is more likely that the detachment of the cortex from the membrane is induced by the unbinding of α-actinin from F-actin rather than by the dissociation of α-actinin from the membrane. However, our assumption in the simulations is expected to have a minor effect on overall results because the dissociation force of ACP from the membrane in the model (∼30 pN) is comparable to that of α-actinin from F-actin. This assumption helps reduce the computational cost.

### 8. Simulation setup

At the beginning of each simulation, an actin network is created via the self-assembly processes of cytoskeletal elements. Actin concentration in all simulations is fixed at *C*_A_ = 10 μM, which is calculated using the entire volume within the membrane. The molar ratios of ACPs (*R*_X_ = *C*_X_ /*C*_A_) and motors (*R*_M_ = *C*_M_/*C*_A_) specified in each simulation determine their amounts. The network is created as either a 2D network right beneath the membrane (i.e., within a space defined by a radial distance from the membrane center between 0.9 × *r*_MB_ and *r*_MB_ where *r*_MB_ is a membrane radius) or a 3D network in an entire space within the membrane (**Fig. 6b**). Note that only 27.1% of the space within the membrane is used for network assembly in case of the 2D network. During the network assembly, the membrane is frozen without a change in its spherical shape, and the network and membrane are coupled via attractive forces acting on ACPs selected at the beginning of simulations. After the network assembly, motors start walking to generate mechanical forces, and the membrane also starts deforming by thermal fluctuation and motor-generated forces.

### 9. Measurements of the formation and size of blebs

To detect bleb formation, the curvature of the membrane is measured every 5 s. To calculate the curvature, we use the discrete mean curvature approximation with consideration of the Voronoi area and common triangular edge length, as explained in detail in a previous study^15^ (**Fig. S6a**). After calculating the curvature, the thresholding technique is used to detect blebs appropriately. To detect blebs accurately and exclude random picks, the threshold curvature was carefully chosen. A region with a high positive curvature surrounded by negative-curvature regions is considered a possible location for bleb formation. Then, distances between the apex of the identified bleb and the edges of the bleb where the curvature flips from positive to negative are measured, and then the average of the distances is calculated as the bleb size. Our algorithm can keep track of more than one bleb emerging in a simulation (**Fig. S6b-d**).

### 10. Measurement of tension acting on the actin network and the membrane

To assess global tension acting on the actin network or the membrane, we consider *xy, yz*, and *zx* cross-sections that include the instantaneous centroid of the membrane for membrane tension or include the centroid of the network for network tension (**Fig. S6e**). All chains of the membrane mesh (for membrane tension) or all segments of F-actin, ACPs, and motors (for network tension) crossing each cross-section are selected. The magnitude of extensional forces acting on selected chains or segments, *F*_s,*i*_, is calculated, which is either positive (= tensile) or negative (= compressive) (**Fig. S6f**). Then, an acute angle between the orientation of chains or segments and a direction normal to the cross-section, *ϕ*_*i*_, is calculated. The sum of *F*_s,*i*_ cos*ϕ*_*i*_ over all chains or segments is considered total tension acting on one cross-section. Then, it is normalized by the contour length of intersection between the network and the cross-section or by the contour length of intersection between the membrane and the cross-section. Global network/membrane tension is calculated by averaging the normalized tension measured on three cross-sections.

To evaluate time evolution of local tension developed near the identified bleb, we first find a time point when the bleb radius reaches 10% of the membrane radius. Then, all F-actin located beneath the bleb and within ∼300 nm from the edges of the bleb is selected. The sum of extensional forces acting on these F-actin is calculated from the end of the network assembly (when motors start walking) till the time point when F-actin is selected.

### 11. Assessment of local areal density of membrane-bound ACPs

First, the surface area of a spherical membrane is divided into quadrilateral and triangular surface elements using the spherical coordinate (**Fig. S6g**). Membrane nodes are allocated to the surface elements based on their positions. Since the distance between neighboring membrane nodes is typically smaller than the size of one surface element, several membrane nodes are assigned to a single surface element. Then, the local areal density of membrane-bound ACPs on each surface element is calculated by dividing the total number of ACPs coupled to all the membrane nodes allocated to the surface element by the sum of the areas that the membrane nodes occupy. The area occupied by each membrane node is calculated by averaging the instantaneous area of three triangular membrane elements connected to the membrane node.

**Table S1.** List of parameters employed in the model.

| Symbol | Definition | Value |
| --- | --- | --- |
| $l_{0,A}$ | Length of an actin segment | $1.4 \times 10^{-7}$ [m] |
| $d_A$ | Diameter of an actin segment | $7.0 \times 10^{-9}$ [m] (ref. 16) |
| $\theta_{0,A}$ | Bending angle formed by adjacent actin segments | 0 [rad] |
| $\kappa_{s,A}$ | Extensional stiffness of F-actin | $1.69 \times 10^{-2}$ [N/m] |
| $\kappa_{b,A}$ | Bending stiffness of F-actin | $2.64 \times 10^{-19}$ [N·m] (ref. 2) |
| $\kappa_{r,A}$ | Strength of repulsive force between neighboring F-actin | $1.69 \times 10^{-3}$ [N/m] |
| $k_{n,A}$ | Nucleation rate of actin | $1.0 \times 10^{-6}$ [$\mu\text{M}^{-1}\text{s}^{-1}$] |
| $k_{+,A}$ | Polymerization rate of actin at the barbed end | 60 [$\mu\text{M}^{-1}\text{s}^{-1}$] |
| $F_{\text{sev}}$ | A minimum force for F-actin severing | 300 [pN] |
| $l_{0,ACP}$ | Length of an ACP segment | $2 \times 10^{-8}$ [m] (ref. 17) |
| $d_{ACP}$ | Diameter of an ACP segment | $1.0 \times 10^{-8}$ [m] |
| $\theta_{0,ACP}$ | Bending angle formed by two ACP segments | 0 [rad] |
| $\kappa_{s,ACP}$ | Extensional stiffness of ACP | $2.0 \times 10^{-3}$ [N/m] |
| $\kappa_{b,ACP}$ | Bending stiffness of ACP | $1.04 \times 10^{-19}$ [N·m] |
| $k_{+,ACP}$ | Binding rate of an ACP segment | 100 [$\text{s}^{-1}$] |
| $l_{0,M1}$ | Length of a motor backbone segment | $4.2 \times 10^{-8}$ [m] |
| $d_{M1}$ | Diameter of a motor backbone segment | $1.0 \times 10^{-8}$ [m] |
| $\theta_{0,M}$ | Bending angle formed by motor backbone segments | 0 [rad] |
| $\kappa_{s,M1}$ | Extensional stiffness of a motor backbone | $1.69 \times 10^{-2}$ [N/m] |
| $\kappa_{b,M}$ | Bending stiffness of a motor backbone | $5.07 \times 10^{-18}$ [N·m] |
| $l_{0,M2}$ | Equilibrium length 1 of a motor arm | $1.0 \times 10^{-8}$ [m] |
| $l_{0,M3}$ | Equilibrium length 2 of a motor arm | 0 [m] |
| $d_{M2}$ | Diameter of a motor arm | $1.0 \times 10^{-8}$ [m] |
| $\kappa_{s,M2}$ | Extensional stiffness 1 of a motor arm | $1.0 \times 10^{-3}$ [N/m] |
| $\kappa_{s,M3}$ | Extensional stiffness 2 of a motor arm | $1.0 \times 10^{-3}$ [N/m] |
| $N_h$ | Number of heads represented by a motor arm | 4 |
| $N_{\text{arm}}$ | Number of arms in a single motor | 16 |
| $k_{+,M}$ | Binding rate of a motor arm | $40N_h$ [$\text{s}^{-1}$] |
| $l_{0,MB}$ | Length of a membrane mesh chain | $7.0 \times 10^{-8}$ [m] |
| $d_{MB}$ | Thickness of a membrane mesh | $5.0 \times 10^{-8}$ [m] |
| $\theta_{0,MB}$ | Dihedral angle formed by two adjacent mesh elements | 0 [rad] |
| $\kappa_{s,MB}$ | Extensional stiffness of a membrane mesh chain | $1.0 \times 10^{-4}$ [N/m] |
| $\kappa_{b,MB}$ | Bending stiffness of a membrane mesh | $2.4 \times 10^{-19}$ [N·m] |
| $A_{0,MB}$ | Area of one triangular mesh | $2.12 \times 10^{-15}$ [$\text{m}^2$] |
| $\kappa_{A,MB}$ | Strength of areal conservation | $1.0 \times 10^{-3}$ [N/m] |
| $V_{0,MB}$ | Volume within a membrane mesh | $2.68 \times 10^{-16}$ [$\text{m}^3$] |
| $\kappa_{V,MB}$ | Strength of volume conservation | $1.0 \times 10^3$ [N/$\text{m}^2$] |
| $\kappa_{a,MB}$ | Strength of an attractive force between a mesh and ACPs | $1.0 \times 10^{-3}$ [N/m] |
| $\kappa_{r,MB}$ | Strength of a repulsive force acting between all cytoskeletal elements and a mesh and between membrane mesh elements | $5.1 \times 10^{-3}$ [N/m] |
| $\Delta t$ | Time step | $1.15 \times 10^{-4}$ [s] |
| $\mu$ | Viscosity of a medium | 8.6 [Pa·s] |
| $k_B T$ | Thermal energy | $4.142 \times 10^{-21}$ [J] |

**Table S2.** List of parameter values used for adopting “parallel cluster model”. Note that we used slightly different values for *F*_0,M_, *s*, and *k*_m_ from those in the literatures (refs. 13, 14).

| Symbol | Definition | Value |
| --- | --- | --- |
| $k_{01,M}$ | A rate from unbound to weakly bound state | 40 [$s^{-1}$] |
| $k_{10,M}$ | A rate from weakly bound to unbound state | 2 [$s^{-1}$] |
| $k_{12,M}$ | A rate from weakly bound to post-power-stroke state | 1000 [$s^{-1}$] |
| $k_{21,M}$ | A rate from post-power-stroke to weakly bound state | 1000 [$s^{-1}$] |
| $k_{20,M}$ | A rate from post-power-stroke to unbound state | 20 [$s^{-1}$] |
| $F_{0,M}$ | Constant for force dependence | $5.04 \times 10^{-12}$ [N] |
| $E_{pp}$ | Free energy bias toward the post-power-stroke state | $-60 \times 10^{-21}$ [J] |
| $E_{ext}$ | External energy contribution | 0 [J] |
| $s$ | Step size | $7 \times 10^{-9}$ [m] |
| $k_m$ | Spring constant of the neck linkers | $1.0 \times 10^{-3}$ [N/m] ($=\kappa_{s,M3}$) |

## MOVIE CAPTIONS

**Movie S1. Liposome showing bleb formation induced by the contraction of a volume-spanning actin network (3D network)**. The liposome contains 10 μM actin, 2 μM His-α-actinin, 1 μM SMM (10% Alexa488-labeled), and 1.4 × 10^−4^ unit ml^-1^ ZIPK, but did not contain methylcellulose. The images were taken with a confocal microscope. Scale bar, 10 μm.

**Movie S2. Liposome showing local actin network contraction but no detectable deformation (2D network)**. The liposome contains 10 μM actin, 1 μM SMM, 1.4 × 10^−4^ unit ml^-1^ ZIPK, and methylcellulose, but did not contain His-α-actinin. The images were taken with an epi-fluorescence microscope. Scale bar, 10 μm.

**Movie S3. Liposome showing weak deformation (2D network)**. The liposome contains 10 μM actin, 0.5 μM His-α-actinin, 1 μM SMM, 1.4 × 10^−4^ unit ml^-1^ ZIPK, and methylcellulose. The images were taken with an epi-fluorescence microscope. Scale bar, 10 μm.

**Movie S4. Liposome forming a single bleb (2D network)**. The liposome contains 10 μM actin, 5 μM His-α-actinin, 1 μM SMM, 1.4 × 10^−4^ unit ml^-1^ ZIPK, and methylcellulose. The images were taken with an epi-fluorescence microscope. Scale bar, 10 μm.

**Movie S5. Liposome forming two blebs (2D network)**. The liposome contains 10 μM actin, 1 μM His-α-actinin, 1 μM SMM, 1.4 × 10^−4^ unit ml^-1^ ZIPK, and methylcellulose. The images were taken with an epi-fluorescence microscope. Scale bar, 10 μm.

**Movie S6. Liposome showing the contraction of an actin network without membrane deformation (2D network)**. The liposome contains 10 μM actin, 2 μM α-actinin, 1 μM SMM, 1.4 × 10^−4^ unit ml^-1^ ZIPK, and methylcellulose. The images were taken with an epi-fluorescence microscope. Scale bar, 10 μm.

**Movie S7. Liposome forming a single bleb initiated by the rupture of the cortical actin network (2D network)**. The liposome contains 10 μM actin, 2 μM His-α-actinin, 1 μM SMM, 1.4 × 10^−4^ unit ml^-1^ ZIPK, and methylcellulose. The images were taken with an epi-fluorescence microscope. Scale bar, 10 μm.

**Movie S8. Liposome forming a single bleb initiated by the detachment of the cortical actin network from the membrane (2D network)**. The liposome contains 10 μM actin, 2 μM His-α-actinin, 1 μM SMM, 1.4 × 10^−4^ unit ml^-1^ ZIPK, and methylcellulose. The detached cortical actin network completed the contraction without any detectable rupture. The images were taken with an epi-fluorescence microscope. Scale bar, 10 μm.

**Movie S9. Liposome forming a single bleb initiated by the detachment of the cortical actin network from the membrane, followed by spontaneous rupture of the contracting actin network (2D network)**. The liposome contains 10 μM actin, 2 μM His-α-actinin, 1 μM SMM, 1.4 × 10^−4^ unit ml^-1^ ZIPK, and methylcellulose. After the detachment of the cortical actin network, the network was spontaneously ruptured (*t* = 31 min). The images were taken with an epi-fluorescence microscope. Scale bar, 10 μm.

**Movie S10. Liposome forming a single bleb with a lower concentration of ZIPK (2D network)**. The liposome contains 10 μM actin, 2 μM His-α-actinin, 1 μM SMM, 0.53 × 10^−4^ unit ml^-1^ ZIPK, and methylcellulose. The images were taken with an epi-fluorescence microscope. Scale bar, 10 μm.

**Movie S11. Liposome showing neither detectable deformation nor actin network contraction (3D network)**. The liposome contains 10 μM actin, 1 μM SMM, and 1.4 × 10^−4^ unit ml^-1^ ZIPK, but did not contain His-α-actinin. The images were taken with an epi-fluorescence microscope. Scale bar, 10 μm.

**Movie S12. Liposome showing weak deformation (3D network)**. The liposome contains 10 μM actin, 0.5 μM His-α-actinin, 1 μM SMM, and 1.4 × 10^−4^ unit ml^-1^ ZIPK. The images were taken with an epi-fluorescence microscope. Scale bar, 10 μm.

**Movie S13. Liposome forming multiple blebs (3D network)**. The liposome contains 10 μM actin, 2 μM His-α-actinin, 1 μM SMM, and 1.4 × 10^−4^ unit ml^-1^ ZIPK. The images were taken with an epi-fluorescence microscope. Scale bar, 10 μm.

**Movie S14. Liposome forming a single bleb (3D network)**. The liposome contains 10 μM actin, 3 μM His-α-actinin, 1 μM SMM, and 1.4 × 10^−4^ unit ml^-1^ ZIPK. The images were taken with an epi-fluorescence microscope. Scale bar, 10 μm.

**Movie S15. Bleb formation in a larger system with 16 μm in diameter (2D network)**. Motor density (*R*_M_) is 0.01, network connectivity (*R*_X_) is 0.01, and actin-membrane coupling strength (*R*_C_) is 0.01.

**Movie S16. Formation of a single bleb under the reference condition (2D network)**. Motor density (*R*_M_) is 0.01, network connectivity (*R*_X_) is 0.08, and actin-membrane coupling strength (*R*_C_) is 0.08. Bleb formation was initiated by the rupture mechanism.

**Movie S17. Successive severing events of F-actin driving the expansion of a rupture formed in the actin network (2D network)**. On the left, local forces acting on the network are visualized via color scaling. On the right, F-actin, actin cross-linking protein, and motor are visualized using different colors.

**Movie S18. No deformation in the absence of actin-membrane coupling (2D network)**. Motor density (*R*_M_) is 0.01, network connectivity (*R*_X_) is 0.08, and actin-membrane coupling strength (*R*_C_) is 0. The network contracted into a smaller cluster without noticeable membrane deformation. Local forces are visualized via color scaling.

**Movie S19. Detachment-induced formation of a single bleb with low actin-membrane coupling strength (2D network)**. Motor density (*R*_M_) is 0.01, network connectivity (*R*_X_) is 0.08, and actin-membrane coupling strength (*R*_C_) is 0.016. Local forces are visualized via color scaling.

**Movie S20. Rupture-induced formation of a single bleb with high actin-membrane coupling strength (2D network)**. Motor density (*R*_M_) is 0.01, network connectivity (*R*_X_) is 0.08, and actin-membrane coupling strength (*R*_C_) is 0.048. Local forces are visualized via color scaling.

**Movie S21. Weak deformation with low network connectivity (2D network)**. Motor density (*R*_M_) is 0.01, network connectivity (*R*_X_) is 0.006, and actin-membrane coupling strength (*R*_C_) is 0.006. Network contraction was observed, but the membrane was not deformed noticeably. Local forces are visualized via color scaling.

**Movie S22. Rupture-induced formation of a single bleb with intermediate network connectivity (2D network)**. Motor density (*R*_M_) is 0.01, network connectivity (*R*_X_) is 0.06, and actin-membrane coupling strength (*R*_C_) is 0.06. The network underwent severe aggregation into a small cluster. Local forces are visualized via color scaling.

**Movie S23. Weak deformation with high network connectivity (2D network)**. Motor density (*R*_M_) is 0.01, network connectivity (*R*_X_) is 0.4, and actin-membrane coupling strength (*R*_C_) is 0.4. The membrane was deformed, but a bleb was not formed. Local forces are visualized via color scaling.

**Movie S24. Formation of multiple blebs (3D network)**. Motor density (*R*_M_) is 0.01, network connectivity (*R*_X_) is 0.08, and actin-membrane coupling strength (*R*_C_) is 0.08. Two blebs were formed by the rupture mechanism. Local forces are visualized via color scaling.

**Movie S25. Distribution, orientation, and force of motors in a 2D network**. Motor density (*R*_M_) is 0.01, network connectivity (*R*_X_) is 0.08, and actin-membrane coupling strength (*R*_C_) is 0.08.

**Movie S26. Distribution, orientation, and force of motors in a 3D network**. Motor density (*R*_M_) is 0.01, network connectivity (*R*_X_) is 0.08, and actin-membrane coupling strength (*R*_C_) is 0.08.

## Notes

### Competing Interest Statement

The authors have declared no competing interest.

